# Early astrocytic dysfunction is associated to mistuned synapses as well as anxiety and depressive-like behavior in the *App^NL-F^* mouse model of Alzheimer’s disease

**DOI:** 10.1101/2023.05.12.540486

**Authors:** Benjamin Portal, Moa Södergren, Teo Parés i Borrell, Romain Giraud, Nicole Metzendorf, Greta Hultqvist, Per Nilsson, Maria Lindskog

**Affiliations:** Department for Medical Cell Biology, Uppsala University, Uppsala, Sweden; Department of Pharmacy, Division of Protein Drug Design, Uppsala University, Uppsala, Sweden; Department of Neurobiology, Care Sciences and Society, Division of Neurogeriatrics, Center for Alzheimer Research, Karolinska Institutet, Stockholm, Sweden

**Keywords:** Alzheimer’s disease, *App* knock-in mice, MAO-B, synapse, LTP, depression

## Abstract

Alzheimer’s disease is the most common neurodegenerative disease and constitute 75% of dementia cases worldwide. Unfortunately, efficient and affordable treatments are still lacking for this mental illness, it is therefore urgent to identify new pharmacological targets. Whereas the late phases of the disease are well described, recent evidence suggest synaptic impairments at a pre-amyloid β (Aβ) plaque stage. Astrocytes are playing a crucial role in the tuning of synaptic transmission and several studies have pointed out severe astrocyte reactivity in Alzheimer’s disease, especially around Aβ plaques. Reactive astrocytes show altered physiology and function, suggesting they could have a role in the early pathophysiology of Alzheimer’s disease. In this study we used the *App^NL-F^* knock-in mouse model of Alzheimer’s disease which carries two disease-causing mutations inserted in the amyloid precursor protein (*App*) gene. This strain does not start to develop Aβ plaques until nine months of age. To better understand early changes in Alzheimer’s disease, we investigated synaptic function, at both neuronal and astrocytic levels, in six months old *App^NL-F^* mice and correlate the synaptic dysfunction with emotional behavior. Electrophysiological recordings in the hippocampus revealed an overall synaptic mistuning at a pre-plaque stage of the pathology, associated to an intact social memory but a stronger depressive-like behavior. Astrocytes displayed a reactive-like morphology and a higher tonic GABA current compared to control mice. Interestingly, we here show that the synaptic impairments in hippocampal slices are partially corrected by a pre-treatment with the monoamine oxidase B (MAO-B) blocker deprenyl or the fast-acting antidepressant ketamine (5mg/kg). Thus, we propose that reactive astrocytes can induce synaptic mistuning early in Alzheimer’s disease, before plaques deposition, and that these changes are associated with emotional symptoms.

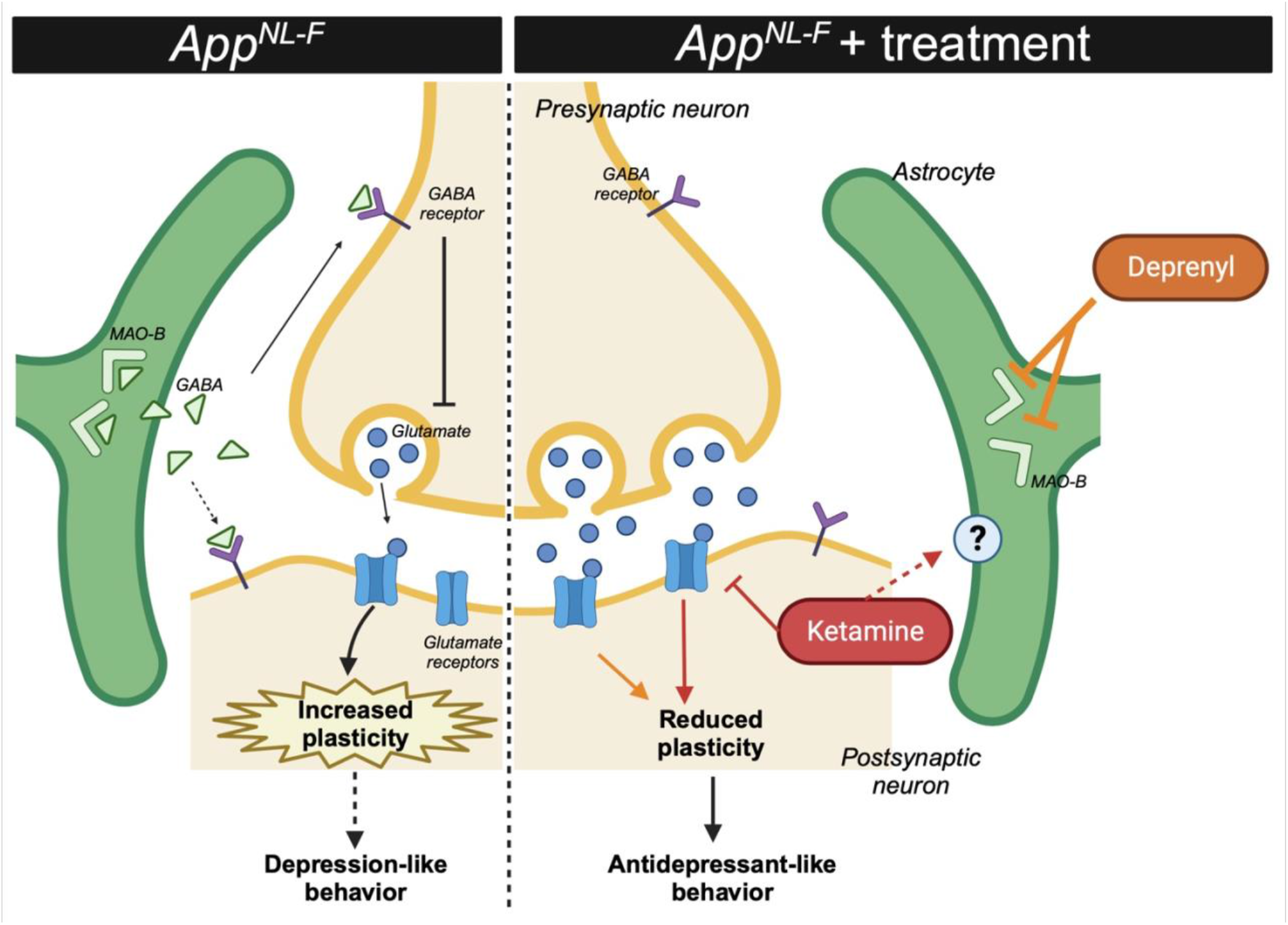
GRAPHICAL ABSTRACT.

## INTRODUCTION

Alzheimer’s disease is a progressive age-related neurodegenerative disorder accounting for 75% of dementia cases worldwide [1]. Patients show significant memory loss [2] together with multiple secondary symptoms such as social isolation [3] and depressive symptoms [4]. In recent years, significant improvements have been done in clinical care of patients and recent anti-Aβ immunotherapies have shown promising results in early phases of the pathology [5]. However, the underlaying mechanisms of this pathology are still unknown. Intracellular tau neurofibrillary tangles [6] and amyloid-beta (Aβ) aggregation [7] are the main molecular hallmarks of the pathology and have been given considerable attention in the attempt to understand the disorder. Extensive cellular rearrangements occur around Aβ plaques, including synaptic degeneration and astrocyte reactivity. In addition, Aβ can take other aggregated forms, including oligomers that are proposed to be the most toxic for neurons [8] and astrocytes [9]. Thus, Aβ oligomers could induce neurobiological changes before plaque deposition.

Early symptoms of Alzheimer’s disease reach beyond memory loss, including sleep disturbance, psychosis, and social isolation [10]. Depressive symptoms are also common in the early stages and depression is a strong risk factor for Alzheimer’s disease. Whether depression is causative, or a prodromal indication is still a question of discussion [11]. Interestingly, antidepressant treatment such as ketamine or monoamine oxidase inhibitors has been shown to reduce cognitive deficits as well as pathology in Alzheimer’s disease [12, 13].

Recent imaging work suggests reduced synaptic density already at early stages of Alzheimer’s disease [14, 15] whereas clinical data show increased functional connectivity and synchronization of neuronal activity early in the disease [16]. These results appear contradictory and we still need to better understand the functional changes of synaptic transmission in early Alzheimer’s disease. In this context, a deficit in neuronal activity set-point such as an imbalance between firing and neuronal plasticity, has been proposed to underlay early development of the pathology [17]. Interestingly, preclinical studies further emphasize the development of early synaptic deficits in Alzheimer’s disease, with demonstration of synaptic impairments at the pre-plaque stage in an *App* knock-in mouse model [18, 19] and impaired synaptic plasticity in a hypercholesterolemia mouse model of Alzheimer’s disease [20].

Reactive astrocytes around plaques are very well described [21] and can be a consequence of Aβ accumulation [22]. More recently, astrocytic reactivity has been demonstrated at an early stage in the progression of Alzheimer’s disease [23, 24] and increased level of the astrocytic protein GFAP has been detected in the cerebrospinal fluid of patients early in the disease [25]. Together with the fact that healthy astrocytes are important for Aβ clearance, thus reactive astrocytes could facilitate Aβ accumulation [26], these findings have led to the proposal of an astrocytic origin of Alzheimer’s [27]. Early astrocyte reactivity in Alzheimer’s disease is compatible with synaptic changes due to the close interaction between astrocytic and synaptic function. Reactive astrocytes affect synapses through several mechanisms [28]. Among other things, reactive astrocytes have been shown to express monoamine oxidase type B (MAO-B) [29] resulting in *de novo* synthesis of GABA and an increased tonic inhibition of adjacent neurons [30].

Many animal models have been engineered in order to understand the mechanisms underlying Alzheimer’s disease, yet none of them recapitulate all the symptoms of the pathology [31]. One common feature in older models is an overproduction of the amyloid precursor protein (App), resulting in an important Aβ accumulation at a young age. The *App^NL-F^* mouse line is a late onset knock-in model of Alzheimer’s disease that is useful to investigate early stages of the pathology. This model carries two disease causing mutations (the Swedish mutation, NL and the Beyreuther/Iberian mutation, F) in the endogenous *App* gene [32] and hence free of App overexpression. The *App^NL-F^* mouse does not develop pathological plaques until nine months of age [33]. While no tau pathology has been unveiled in this model [34], except increased phosphorylation of some sites [35], many molecular changes such as autophagy disturbances [36] and mitochondria dysfunctions [37] were reported both *in vivo* and *in vitro* [38]. Interestingly, early changes in the synapse structure and function has been identified in the cortex of six months old *App^NL-F^* animals [19, 39].

In the present study we used behavioral, electrophysiological and molecular readouts to understand early changes in this *App^NL-F^* knock-in mouse model of Alzheimer’s disease. Our analysis revealed reduced synaptic activity and impaired synaptic plasticity associated to reactive-like astrocytes. These neuronal and astrocytic dysfunctions correlate with increased depressive-like behavior and could be partially corrected by treatment with either the MAO-B inhibitor deprenyl or the fast-acting antidepressant ketamine.

## METHODS

### Animals

*App^NL-F^* mice, model of Alzheimer’s disease, were bred locally at Uppsala University. These mice carry the Swedish (KM670/671NL) and the Beyreuther/Iberian (I716F) mutations in the App gene. Age-matched C57Bl6 mice were bought from Charles River (Germany). Both strains were housed in the same conditions, on a 12:12 light-dark cycle and with access to food and water *ad libitum*. All experiments were conducted in six months old males, where increased insoluble Aß was detected but where Aβ-plaques were absent (fig. S1).

### Drugs

Tetrodotoxin (TTX; Tocris 4368-28-9) was diluted in artificial cerebrospinal fluid (aCSF) and used in patch clamp experiments to measure mini excitatory post-synaptic currents (mEPSC), at a concentration of 1 µM.

NBQX (Tocris 1044), DL-AP5 (Tocris 3693), Picrotoxin (PTX; Tocris 1128) were diluted in aCSF and used in patch clamp experiments to measure tonic GABA current, at a concentration of 10 µM, 50 µM and 100 µM respectively.

(R)-(–)-Deprenyl hydrochloride (deprenyl; Tocris 1095) was diluted in aCSF and was used in the patch clamp and the field recording experiments, at a concentration of 100 µM.

Ketamine (Ketaminol® vet., 100 mg/kg, Intervet) was diluted in vehicle solution (natriumchloride, NaCl 0.9%) and delivered intraperitoneally (i.p.; 5 mg/kg) twenty-four hours prior to experiments.

### Behavioral Analysis

Animals were habituated to the experimental room twenty minutes prior to experiments. The luminosity of the room was set on 30 lux and the temperature at 21°C ± 1°C. The tests were performed in the following order, with at least three days between each test.

### Open field test

The animals were free to explore a spatial-cue-free square arena (50cm x 50cm) for ten minutes. The central zone and peripheral zone were defined in the Ethovision software (Noldus tech, The Netherlands). The area of the central zone was defined as 50% of the area of the arena. The total travelled distance, the time spent in the central area, as well as the number of entries in the central zone, were automatically analyzed by the software.

### Five-trials social memory test

The mice were habituated to an empty square arena (50 cm x 50 cm) for ten minutes, followed by five minutes exploration of the same arena with an empty removable cage (8cm x 8 cm x 9 cm). An unknown mouse (same genotype, same fur color, same age, and same gender as the test mouse) was placed for five minutes in the removable cage for four consecutive trials, separated by ten minutes. One hour after the fourth trial, a new unknown mouse was placed in the cage for five minutes. Social interactions were manually scored.

### Olfactory habituation/dishabituation test

The mice were isolated in individual standard cages in which they performed the whole experiment and habituated for at least twenty minutes. Cotton swabs soaked with non-social odors (melon and banana) or social odors (one form their home cage and one from a non-familiar cage) were presented to the animal for two minutes. Each odor was presented three times.

### Elevated plus maze

The maze was composed of two open arms, two closed arms, linked together at a central platform and positioned 50 cm above ground. Mice were placed on the central platform facing one open arm and were let to explore for five minutes. The number of entries and the time spent in the open arms were manually scored. An animal was considered “in the arm” when the four paws crossed the virtual line between the central platform and the considered arm.

### Forced swim test

The mice were placed in a cylinder (30 cm high, ∅ 18 cm) filled with 25 ± 1°C water to a height of 17 cm where no part of the animal could touch the bottom. The mice were left in the cylinder for six minutes, and the time spend immobile was quantified during the last four minutes. Latency to first immobility and the total time of immobility were manually scored. An animal was considered as immobile when it was floating and none of the paws nor the tail was moving.

### Emotionality z-score

Z-normalization was used in this study as a complementary measurement for emotionality-related behavior, obtain from different paradigms [40]. Raw behavioral data (parameters of interest) were normalized to the control group using the following equation *z* = (*X* − μ)/σ with X being the individual value for the considered parameter, µ being the mean of the control group and σ being the standard deviation of the control group. Each parameter of interest gave a parameter z-score (zSC_param), adjusted so that an increased score reflects an increased emotionality. For each test, all parameter scores were averaged into a test score (zSC_test), and eventually averaged to give an individual emotionality z-score.

More details about the exact list of parameters of interest and the mathematical method are available in table S1 and table S2.

### Electrophysiology

#### Brain slicing

For the preparation of acute slices, mice were anesthetized with isofluorane and decapitated soon after the disappearance of corneal reflexes. 300 µm thick horizontal sections were prepared using a Leica VT1200 vibrating microtome (Leica Microsystems, Nussloch, Germany) in dissection solution containing 250 mM sucrose, 2.5 µM KCl, 1.4 mM NaH_2_PO_4_, 26 mM NaHCO_3_, 10 mM glucose, 1 mM CaCl_2_ and 4 mM MgCl_2,_ and bubbled with carbogen gas (5% CO_2_, 95% O_2_). Hippocampi were dissected and slices were placed in a recovery chamber filled with aCSF containing (in mM): 130 NaCl, 3.5 KCl, 1.25 NaH_2_PO_4_, 24 NaHCO_3_, 10 glucose, 2 CaCl_2_ and 1.3 MgCl_2_ and bubbled with carbogen gas.

#### Patch clamp recording

After a recovery period of at least two hours, slices were transferred in a submerged recording chamber with a perfusion rate of 2-3 mL per min with standard aCSF, tempered at 32 ± 1°C and bubbled with carbogen gas. Borosilicate glass pipettes with a tip resistance of 4-5 MΩ were used for patching neurons. The glass pipettes were filled with a solution containing (in mM) 110 K-gluconate, 10 KCl, 4 Mg-ATP, 10 Na_2_-phosphocreatine, 0.3 Na-GTP, 10 4-(2-hydroxyethyl)piperzine-1-ethanesulfonic acid (HEPES) and 0.2 ethylene glycol tetraacetic acid (EGTA) (pH 7.2-7.4; 270-290 mOsm). Pyramidal neurons were identified by shape and localization in stratum radiatum of area CA1 of the hippocampus. Spontaneous excitatory post-synaptic currents (sEPSC) were recorded in absence of any drugs, whereas mEPSC were recorded in the presence of 1 µM TTX. To block the monoamine oxidase B (MAO-B), deprenyl (100 µM) was added in the recovery chamber and all the recording were done under tonic deprenyl treatment.

The access resistance was monitored throughout the recordings and data were included only for stable values (< 30% variation). The signal was acquired using an Ag/AgCl electrode connected to a Multiclamp 700B amplifier, digitized with Digidata 1440A and handled with the Clampex software (v. 10.0; Molecular Devices). Traces were analyzed using the Easy electrophysiology software (v. 2.4). The event detection was based on a template fitting method. Probe recordings (three recordings of one minute each) from C57Bl6 wild type acute hippocampus slices were used to generate and refine a template from representative events corresponding to an average of ten to twenty detected EPSC. Event detection in *App^NL-F^* and C57Bl6 controls acute slices was then run on a semi-automatic method: each event detected fits the previously generated template. A minimum amplitude threshold of 10 pA was applied: all events that were detected between 0 pA and 10 pA were discarded. Each recording was analyzed on a period of 1 minute.

GABA currents [41, 42] were recorded in the presence of NBQX (10 μM) and DL-AP5 (50 μM). After recording a stable baseline, picrotoxin (PTX; 100 μM) was added to block GABA_A_ receptors. Tonic GABA current was calculated as the difference in mean holding current for one minute after achieving a stable shift in baseline with PTX compared to before PTX.

#### Field recording

After a recovery period of at least two hours, slices were transferred in a submerged recording chamber with a perfusion rate of 2-3 ml per minute with either standard aCSF or deprenyl (100 µM)-containing aCSF, tempered at 32±1 °C and bubbled with carbogen gas. An extracellular borosilicate recording pipette filled with aCSF was placed in the stratum radiatum of area CA1 and field Excitatory Post-Synaptic Potentials (fEPSP) were evoked by electrical stimulation of the Schaffer collaterals using a bipolar concentric electrode (FHC Inc., Bowdoin, ME), connected to an isolated stimulator (Digitimer Ltd., Welwyn Garden City, UK). Recordings were performed with the stimulus intensity to elicit 30-40% of the maximal response and individual synaptic responses were evoked at 0.05 Hz (every 20 seconds). The acquired signal was amplified and filtered at 2 kHz (low-pass filter) using an extracellular amplifier (EXT-02F, NPI Electronic, Tamm, Germany). Data were collected and analyzed using a Digidata 1440A, Axoscope and Clampfit softwares (Molecular Devices, San Jose, CA). Responses were quantified by determining the slope of the linear rising phase of the fEPSP (from 10% to 70% of the peak amplitude). The response was normalized to the average baseline measured in the last ten minutes prior to the start of the experiment.

To study long term synaptic plasticity, we used ten theta-bursts (θ-burst) separated with 200 ms. Each θ-burst is four pulses at 100 Hz, a stimulation protocol tested not to be enough to trigger Long Term Potentiation (LTP) in slices from C57Bl6 mice.

### Immunohistochemistry

#### Perfusion and brain extraction

Mice were deeply anesthetized with a mixture of ketamine (100 mg/kg) and xylazine (16 mg/kg) i.p. In absence of corneal reflexes, mice were transcardially perfused with NaCl 0.9% followed by a 4% PFA solution. The brain was extracted and post-fixed in a 4% PFA solution for forty-eight hours, washed in PBS for twenty-four hours and placed in a 30% sucrose + 1% natrium azide solution until further use.

#### Slicing and staining

The brains were sliced in 140 µm thick section using a cryostat (Thermo Scientific cryostar NX70). Free-floating slices were treated with a peroxide solution (10% methanol and 10% H_2_O_2_ in PBS) for twenty minutes followed by antigen retrieval solution (Tris 10 mM EDTA, 10% Tween-20) at eighty degrees Celsius. After blocking steps with normal donkey serum, slices were incubated with primary Glial Fibrillary Acidic Protein (GFAP, Synaptic system 173-004; 1:500) and primary connexin 43 (Cx43, Invitrogen 71-0700; 1:500) for seventy-two hours. Slices were incubated with the appropriate secondary antibodies for two hours, DAPI (1:5000) was added to counterstain nuclei. Slices were then mounted on non-coated microscope glass coverslips using the fluoromount mounting medium (Invitrogen, 00-4958-02).

One z-stacks (0.25 µm step size) per slice (two slices per animal), were obtained from the CA1 region of the hippocampus using a confocal microscope (Zeiss LSM700) with a 63x oil objective.

#### Image analysis

The z-stacks were deconvoluted using the Hyugens software (Scientific Volume Imaging) and noise-saturated pixels were removed. GFAP/DAPI images were segmented thanks to a pixel classification protocol using the Ilastik software [43]. Using Image J (NIH), centroids of the nuclei were determined, DAPI/GFAP images were merged and each astrocyte were manually separated from the astrocytic syncytium thanks to the Cx43 staining giving the precise location of each connexon. Every astrocyte touching at least one border of the z-stack was automatically removed in order to exclude truncated astrocytes. The center of the sholl analysis was defined by the centroid of the nuclei. Branch counting started at a radius 7 of µm and ended at 80 µm from the center, with a stepsize of 2 µm. Subsequent data (i.e., sholl analysis, volume occupied by the cell, number of processes) were automatically generated thanks to the Image J software using a custom macro. Detailed workflow is available in the fig. S2.

### Statistical analysis

IBM SPSS Statistics (v. 28 IBM corp) was used to perform the statistical analysis. For each experiment, a Shapiro-Wilk test for normality and Levene’s test for homogeneity of variances were used to determine the subsequent test to compared means. For all the tests, 0.05 was set as the significance threshold. All results are shown as mean ± S.E.M.

A summary of statistics is available in table S3.

## RESULTS

### Astrocytes display reactive-like phenotype in six months old App^NL-F^ mice

Reactive astrocytes are characterized by a swollen soma and increased branching. Morphology analysis of GFAP stained astrocytes in brain sections from six months old mice revealed a reactive-like morphology in *App^NL-F^* mice compared to C57Bl6 mice (fig. 1A-D). The morphological complexity was quantified as an increased number of branches, counted every 5 µm from the soma (Two-way ANOVA with distance and genotype as the main factors. Distance: F_(8,230)_=98.167, p<0.001; Genotype: F_(1,40)_=26.845; p<0.001; Interaction: F_(8,230)_=19.927, p<0.001). The difference was particularly prominent between 5 and 30 µm from the soma (fig. 1B). Moreover, the total number of branches was higher in *App^NL-F^* compared to C57Bl6 mice (182.6 ± 20.25 vs 45.80 ± 6.759; p<0.001; fig. 1C) as was the volume occupied by the astrocyte (C57Bl6: 2396 ± 486.1 µm^3^ vs. *App^NL-F^:* 10133 ± 1568 µm^3^, p<0.01; fig. 1D). In addition to morphological changes, reactive astrocytes have been shown to synthesize and release GABA [30] and we thus recorded tonic GABA currents in pyramidal neurons in area CA1 (fig. 1E). Indeed, tonic GABA currents were significantly higher in *App^NL-F^* mice compared to C57Bl6 (103.8 ± 22.63 pA vs. 39.83 ± 9.057 pA, p<0.05; fig. 1F).

**Figure 1:**
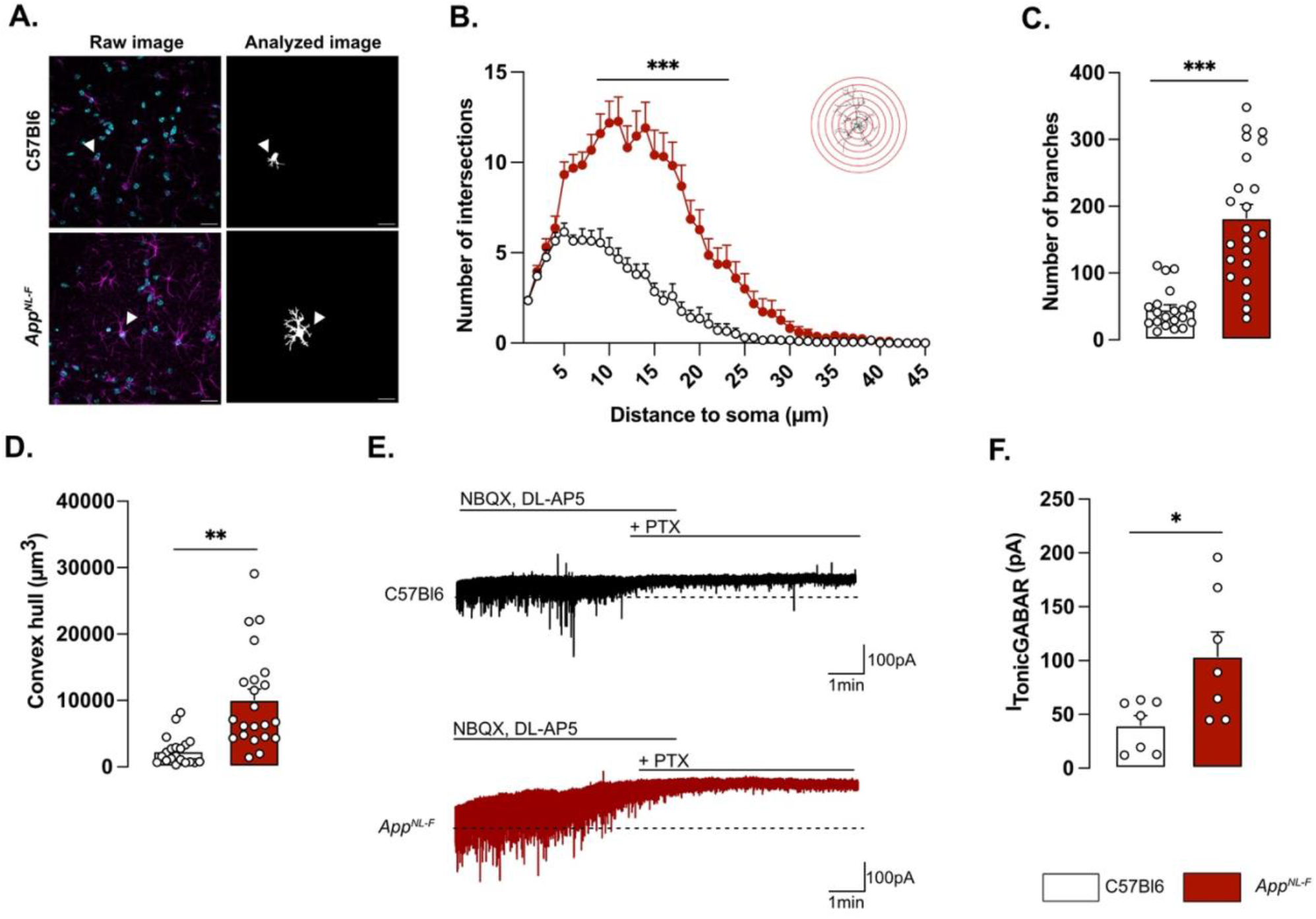
Reactive-like astrocytes in six months old App^NL-F^ mice. Confocal z-stack images of immunohistochemical labelled GFAP-positive astrocytes in area CA1 of the hippocampus were used to analyze astrocytes morphology. Magenta: GFAP, Cyan: DAPI (A. left). Images were deconvoluted and astrocytes were segmented and isolated based on connexin staining. Representative of analyzed image are displayed in A. right panel. White arrows show the representative astrocyte. Scale bare for both view: 20µm. Sholl analysis revealed an increased number of intersections from 5 to 30 µm around the soma (B.) Mann-Whitney test: ***, p<0.001 significant from C57Bl6. The number of branches (C.) and the average volume occupied by the astrocyte (D.) were also increased in App^NL-F^ compared to C57Bl6 mice (Mann-Whitney test: **, p<0.01; ***, p<0.001 significantly different as shown). For all measurements, C57Bl6: n=20 cells in three animals, App^NL-F^: n=22 cells in three animals. Tonic GABA currents were recorded in pyramidal cells in the presence of NBQX and DL-AP5. Representative traces with baseline before blocking GABA is shown as a dashed line (E.). Average tonic GABA currents are significantly higher in App^NL-F^ mice compared to C57Bl6 (F.) Mann-Whitney test: *, p<0.05 significantly different as shown. C57Bl6: n=7 cells recorded in five animals; App^NL-F^: n=7 cells recorded in four animals.

### Impaired synaptic transmission and plasticity in App^NL-F^ mice

Whole-cell patch clamp recordings from pyramidal neurons of area CA1 showed that resting membrane potential was significantly depolarized in *App^NL-F^* mice compared to C57Bl6 mice (fig. 2A; –51.74 ± 1.32 mV vs –56.18 ± 1.38 mV; p<0.05). Spontaneous synaptic activity in active hippocampal network (sEPSC) was not different between the two strains in neither frequency amplitude, decay or rise time of sEPSC (fig. S3A-D). However, when action potentials were blocked with the sodium channel blocker TTX (1 µM) to record spontaneous release of individual synaptic vesicles (mEPSC), a reduced frequency of the mEPSC (fig. 2B) was observed in *App^NL-F^* mice compared to C57Bl6 mice (0.27 ± 0.04 Hz vs. 0.56 ± 0.08 Hz; p<0.01), whereas the amplitude of mEPSCs remained unchanged (C57Bl6: –21.71 ± 1.48 pA, *App^NL-F^*: –19.65 ± 1.43 pA, fig. 2C). There were no differences in the decay or rise time of mEPSC (fig. S3E, F respectively).

**Figure 2:**
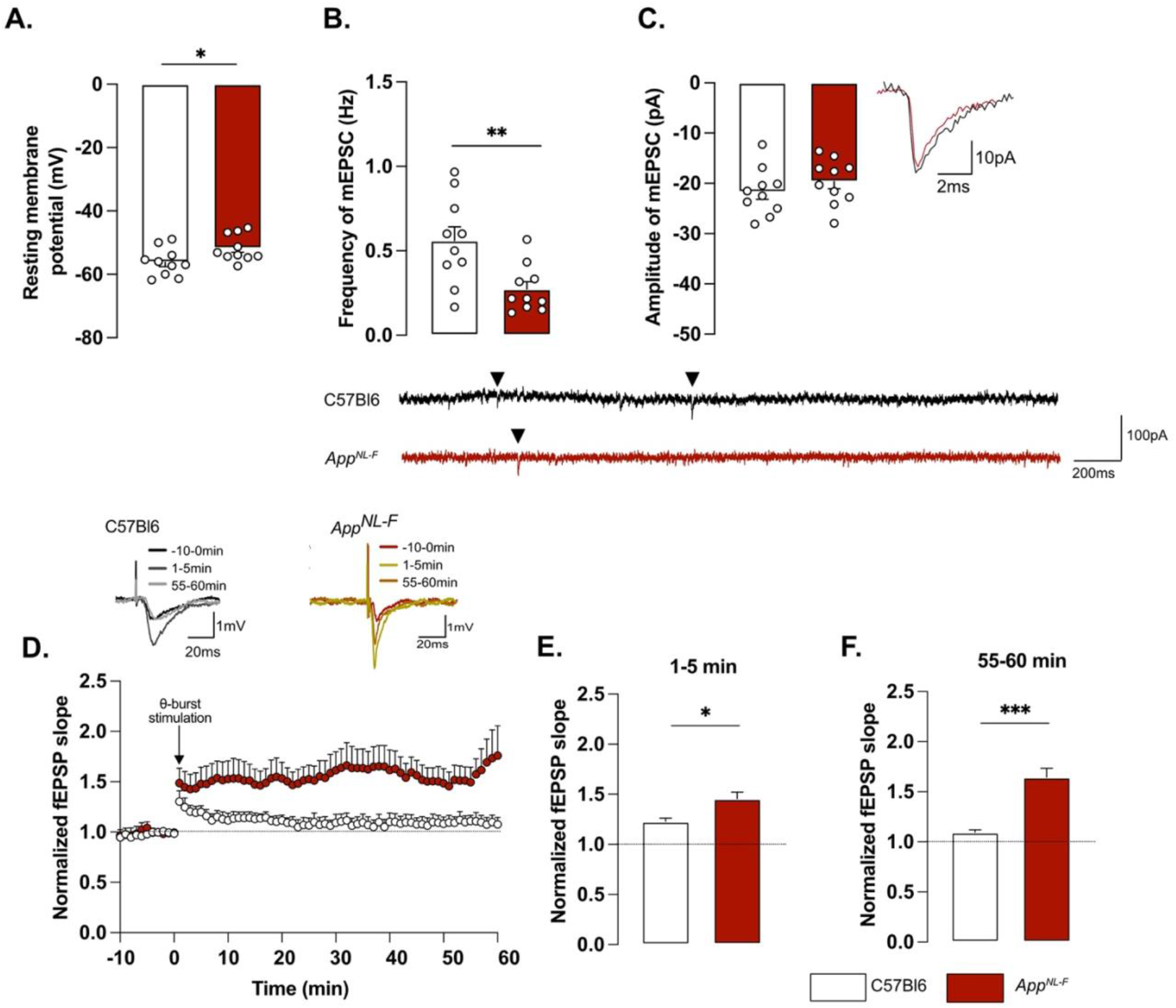
Mistuned synapses in six months old App^NL-F^ mice. Average resting membrane potential of pyramidal neurons of area CA1 of the hippocampus is significantly higher in App^NL-F^ mice compared to C57Bl6 mice (A.). Recordings of mEPSC in the same cells show reduced frequency of events (B.) in App^NL-F^ compared to C57Bl6 mice, with no changes in amplitude (C.). Representative traces for mEPSC are shown under graphs. Student t-test, **, p<0.01 statistically significant as shown. For all variables, C57Bl6: n=10 cells recorded in seven animals, App^NL-F^: n=10 cells recorded in seven animals. fEPSP were recorded after Schaffer collaterals stimulation in the area CA3. We recorded the response of post-synaptic neurons in area CA1. A subthreshold θ-burst stimulation was applied and fEPSP magnitude was monitored for an hour. The fEPSP slope was normalized to the average baseline value and displayed as the average per minute (D.). Representative traces are shown on top. The θ-burst induced a significant potentiation in both AppNL-F and C57Bl6 mice at 0-5 minutes after stimulation (E.). However, at 55-60 minutes, the fEPSP magnitude is almost back to baseline in C57Bl6 mice and is significantly increased in App^NL-F^ mice (F.). Student t-test: *, p<0.05, ***, p<0.001 statistically significant as shown. C57Bl6: n=6 recordings in three animals, App^NL-F^: n=9 recordings in five animals.

Changes in basic synaptic transmission can entail changes in synaptic plasticity, previously described as synaptic mistuning [44]. To investigate the state of plasticity in *App^NL-F^*, the possibility to undergo long-term potentiation (LTP) was examined, using a subthreshold stimulation protocol that did not evoke LTP in C57Bl6 mice. Ten theta-bursts (θ-burst) stimulation of Schaffer collaterals induced an immediate potentiation of synaptic strength, as measured as the increased slope of the evoked fEPSP, in both genotypes (p<0.001; fig. 2D, E). However, one hour after stimulation, the fEPSP had progressively returned to baseline in control animals, whereas the magnitude of the fEPSP was still significantly increased in *App^NL-F^* mice (p<0.001 different from the baseline; fig. 2D, F). Thus, already at six months, *App^NL-F^* mice have changes in synaptic function, detected as reduced frequency of synaptic events and a reduced threshold for potentiation.

### MAO-B blocking restores synaptic transmission and plasticity in App^NL-F^ mice

To examine the relationship between astrocytic GABA and synaptic function, the astrocytic GABA-synthesizing enzyme MAO-B was blocked by pre-incubating the slices with deprenyl (100 µM). Blocking MAO-B caused an increase in the resting membrane potential in deprenyl treated slices from *App^NL-F^*mice compared to non-treated slices (–54.84 ± 1.994 mV vs. 46.30 ± 3.251 mV, p<0.05, fig. 3A.). Moreover, deprenyl increased the frequency of mEPSC events from 0.05 ± 0.008 Hz, in non-treated slices to 0.26 ± 0.03 Hz in deprenyl treated slices (p<0.01, fig. 3B.) and the amplitude from –15.23 ± 2.10 pA to –23.06 ± 1.79 pA (p<0.05, fig. 3C). Deprenyl did not affect rise time or decay time of either sEPSC or mEPSC (fig. S3I-L). For spontaneously evoked synaptic events, deprenyl treatment significantly increased the amplitude of sEPSC but not the frequency (fig. S3G, H).

**Figure 3:**
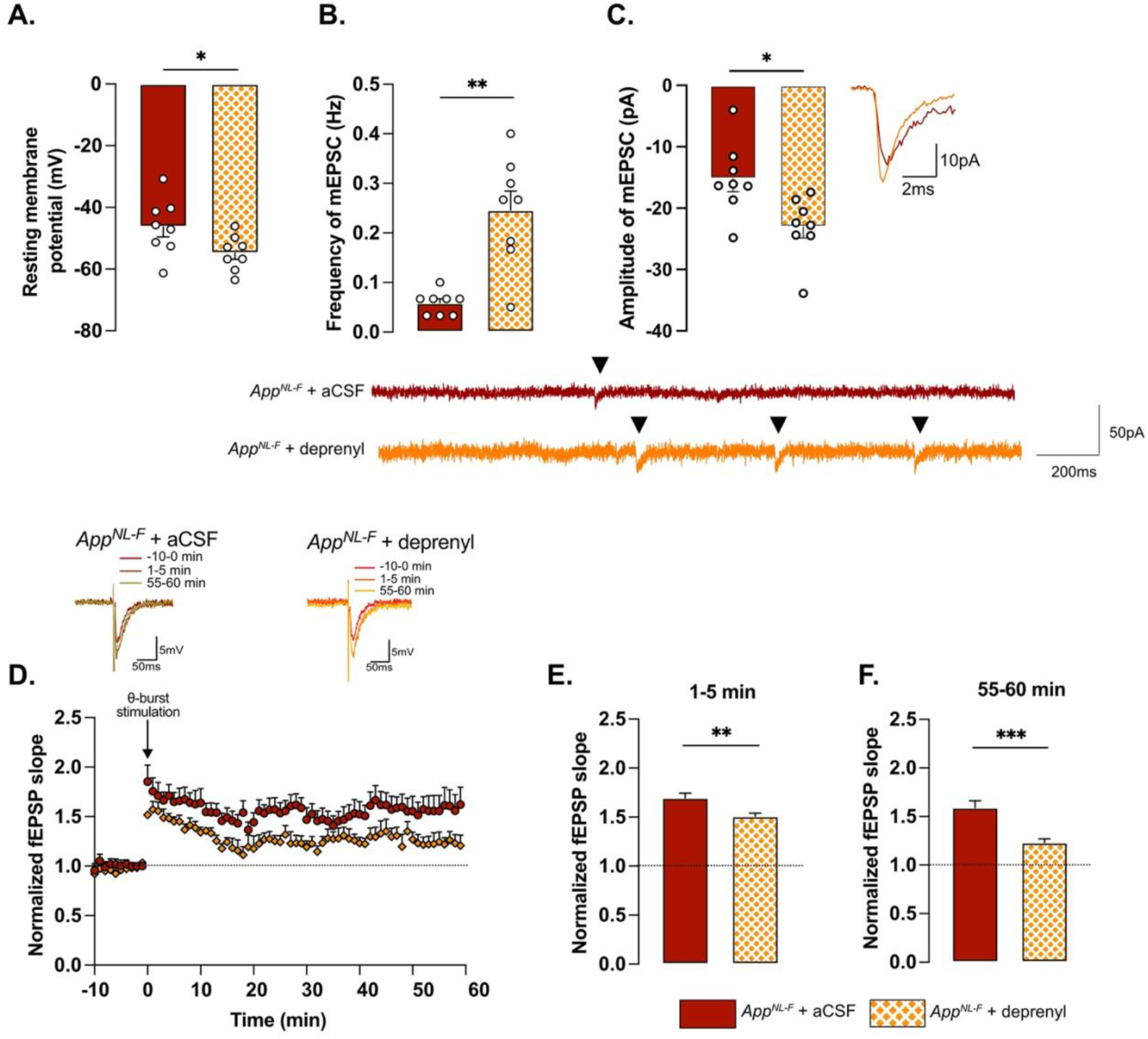
Retuned synapses after MAO-B blocking by deprenyl in six months old App^NL-F^ mice. Pre-treatment of hippocampal slices with the MAO-B blocker deprenyl significantly decreases resting membrane potential in pyramidal cells in App^NL-F^ mice (A.). Blocking MAO-B increased both frequency (B.) and Amplitude (C.) of mEPSC. Representative traces are shown under graphs. Student t-test and Mann-Whitney test: **, p<0.01; ***, p<0.001 statistically significant as shown. App^NL-F^ + aCSF: n=8 cells recorded in four animals; App^NL-F^ + deprenyl: n=8 cells recorded in four animals. The subthreshold θ-burst stimulation induced long term potentiation in both treated and untreated slices from App^NL-F^ mice (D.). Representative traces are shown on top. When we compared the average fEPSP magnitude at 0-5 minutes and 55-60 minutes after stimulation, the potentiation was significantly lower after deprenyl pre-treatment (E., F.). Mann-Whitney test: **, p<0.01, ***, p<0.001 statistically significant as shown. App^NL-F^ + aCSF: n=6 recordings in four animals, App^NL-F^ + deprenyl: n=5 recordings in three animals.

Pre-treatment with deprenyl also had an effect on synaptic plasticity. Unexpectedly, synaptic potentiation was reduced in slices from *App^NL-F^*mice that had been pretreated with deprenyl compared to non-treated slices (fig. 3D). Deprenyl reduced fEPSP magnitude both at 1-5 minutes (*App^NL-F^* + aCSF: 1.70 ± 0.05, *App^NL-F^* + deprenyl: 1.51 ± 0.03) and 55-60 minutes (*App^NL-F^* + aCSF: 1.519 ± 0.07, *App^NL-F^* + deprenyl: 1.23 ± 0.04) after the same sub-threshold stimulation as used above (fig. 3E, F).

Taken together, our results show that already at a pre-plaque stage *App^NL-F^* mice have impaired synaptic function as well as reactive-like astrocytes. Blocking astrocytic GABA synthesis with deprenyl restores synaptic function in the *App^NL-F^* mice.

### App^NL-F^ mice show a lack of motivation and a mild depressive-like behavior

Deficits in plasticity is typically linked to memory problems. However, despite deficits in plasticity at six months, *App^NL-F^*mice have no spatial memory impairment at this age [45]. Other forms of memory have not been explored, prompting us to investigate the possible effect on social memory. In the five-trials social memory test the social interactions with an unknown animal in four consecutive trials is used to evaluate social memory whereas the interaction with a new animal on the fifth trial is used to assess the motivation (fig. 4A). A similar decrease in social interactions was observed over the four first trials in both *App^NL-F^* and C57Bl6 animals (fig. 4B), suggesting no social memory impairments at six months of age. However, on the fifth trial the social interactions with a new animal were increased in control mice (8.93 ± 2.98 seconds on trial 4 vs. 28.41 ± 2.98 seconds on trial 5; p<0.05) whereas no change was observed in the fifth trial compared to the fourth in *App^NL-F^*mice (10.74 ± 1.95 seconds on trial 4 vs. 12.92 ± 4.19 seconds on trial 5). The latency to first interaction was similar in both groups during the first trial, (C57Bl6: 26.32 ± 13.31 seconds, *App^NL-F^*: 11.69 ± 6.903 seconds), however it was significantly increased in *App^NL-F^* mice compared to C57Bl6 mice on the fifth trial (132.01 ± 41.24 seconds vs. 9.73 ± 5.13 seconds; p<0.01; fig. 4C). Together, these results show an intact social memory in *App^NL-F^* mice at six months, and suggest a slight loss of motivation, evident at the fifth trial.

**Figure 4:**
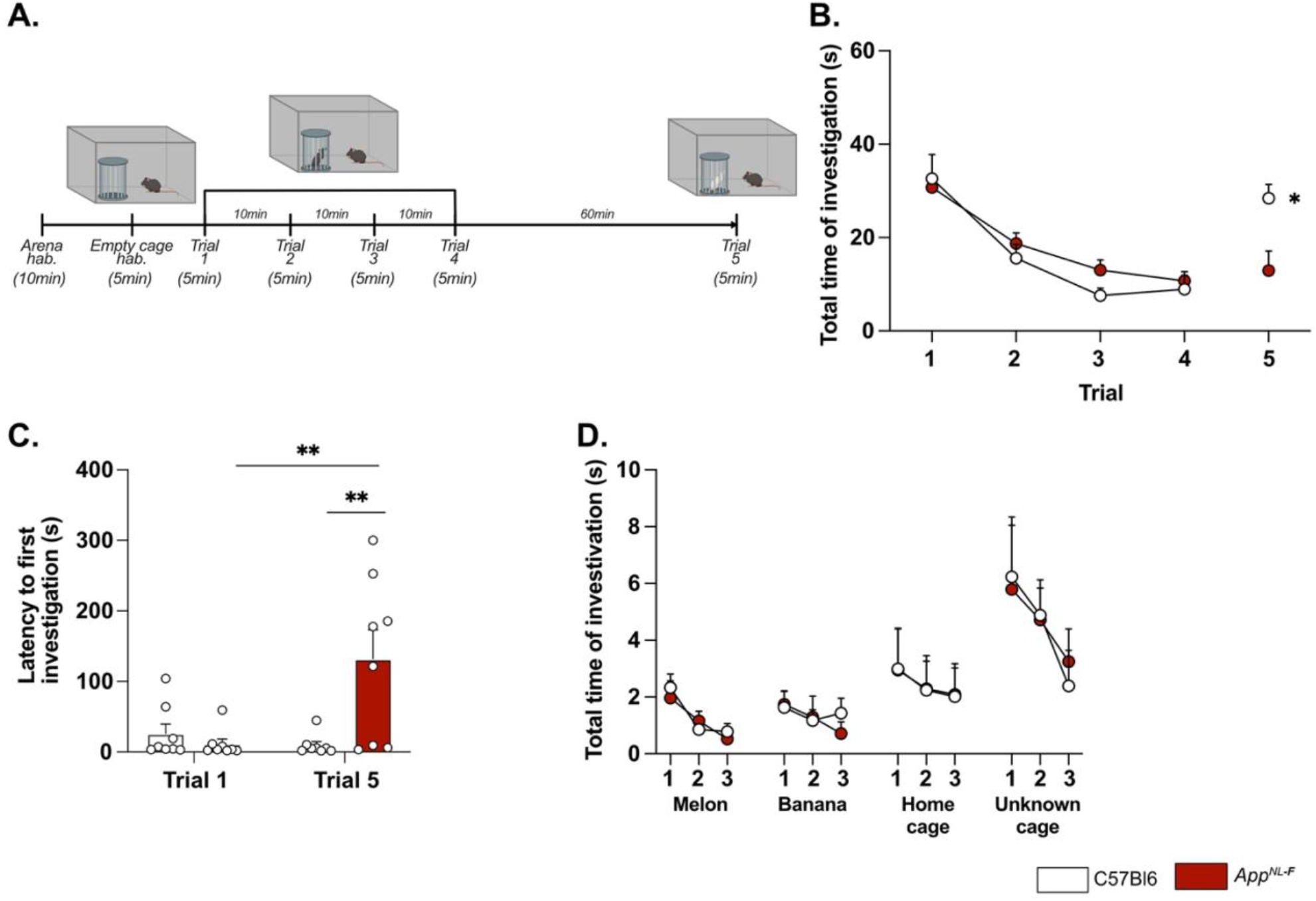
AppNL^-F^ mice shown no social memory impairments but a lack of motivation. In the five-trials social memory test (A.), App^NL-F^ mice show no memory deficits as seen by the progressive decrease in social interactions over four consecutive trials (B.). On the fifth trial, when a new animal is presented, C57Bl6 mice regain interest and the time of interaction is increased, whereas the time of interaction is not different in the fifth trial versus the fourth one in App^NL-F^ mice. Two-way ANOVA for repeated measured with the genotype and the trial as the main factors. Bonferroni multiple comparison *, p<0.05 statistically different trial 4 vs. trial 5. Latency to first investigation (C.) is similar between genotypes in the first trial. However, it is significantly increased in App^NL-F^ on the fifth trial compared to C57Bl6 and to App^NL-F^ in trial 1. Two-way ANOVA for repeated measured with the genotype and the trial as the main factors. Bonferroni multiple comparison; Bonferroni multiple comparison: **, p<0.01: statistically different as shown. C57Bl6: n=8, App^NL-F^: n=8. In the olfactory habituation/dishabituation test (D.), we presented four different odors for three times two minutes each. The time of interaction with the probe was similar in C57Bl6 and App^NL-F^ at all timepoints and for all odors. C57Bl6: n=9, App^NL-F^: n=10.

To rule out potential olfactory dysfunctions, which could interfere with the social memory tests, we performed an olfactory habituation/dishabituation test ([46]; fig. 4D). Four odors were tested and no significant differences between odor recognition was observed between C57Bl6 and *App^NL-F^* mice. Both strains spent more times interacting with social odors, especially from an unknown cage, showing the absence of olfactory problem as well as a capacity of *App^NL-F^* mice to discriminate between social and non-social odors.

A higher level of apathy or lack of motivation in six months old *App^NL-F^*mice is consistent with clinical studies on Alzheimer’s patients [47] and prompted further research of emotional symptoms. In the open field test *App^NL-F^* animals had a significant reduction in the of the travelled distance compared to C57Bl6 (3543 ± 61.1 cm vs. 5663 ± 421.1 cm; p<0.001; fig. 5A). The number of entries in the central area was significantly lower in the *App^NL-F^* mice compared to C57Bl6 mice. (6.50 ± 1 entries vs. 30.75 ± 4 entries; p<0.01; fig. 5B) whereas the time spent in the central zone was not significantly different (C57Bl6: 81.16 ± 6.23 seconds, *App^NL-F^*: 82.30 ± 7.22 seconds; fig. 5C). In the elevated plus maze the number of entries in the open arms (anxiogenic area) was significantly reduced in *App^NL-F^* mice compared to C57Bl6 (3.25 ± 1 entries vs. 6.12 ± 1 entries; p<0.05; fig. 5D) whereas the total time spent in this area was not different between the strains (C57Bl6: 38.38 ± 5.68 seconds, *App^NL-F^*: 26.63 ± 5.76 seconds; fig. 5E).

**Figure 5:**
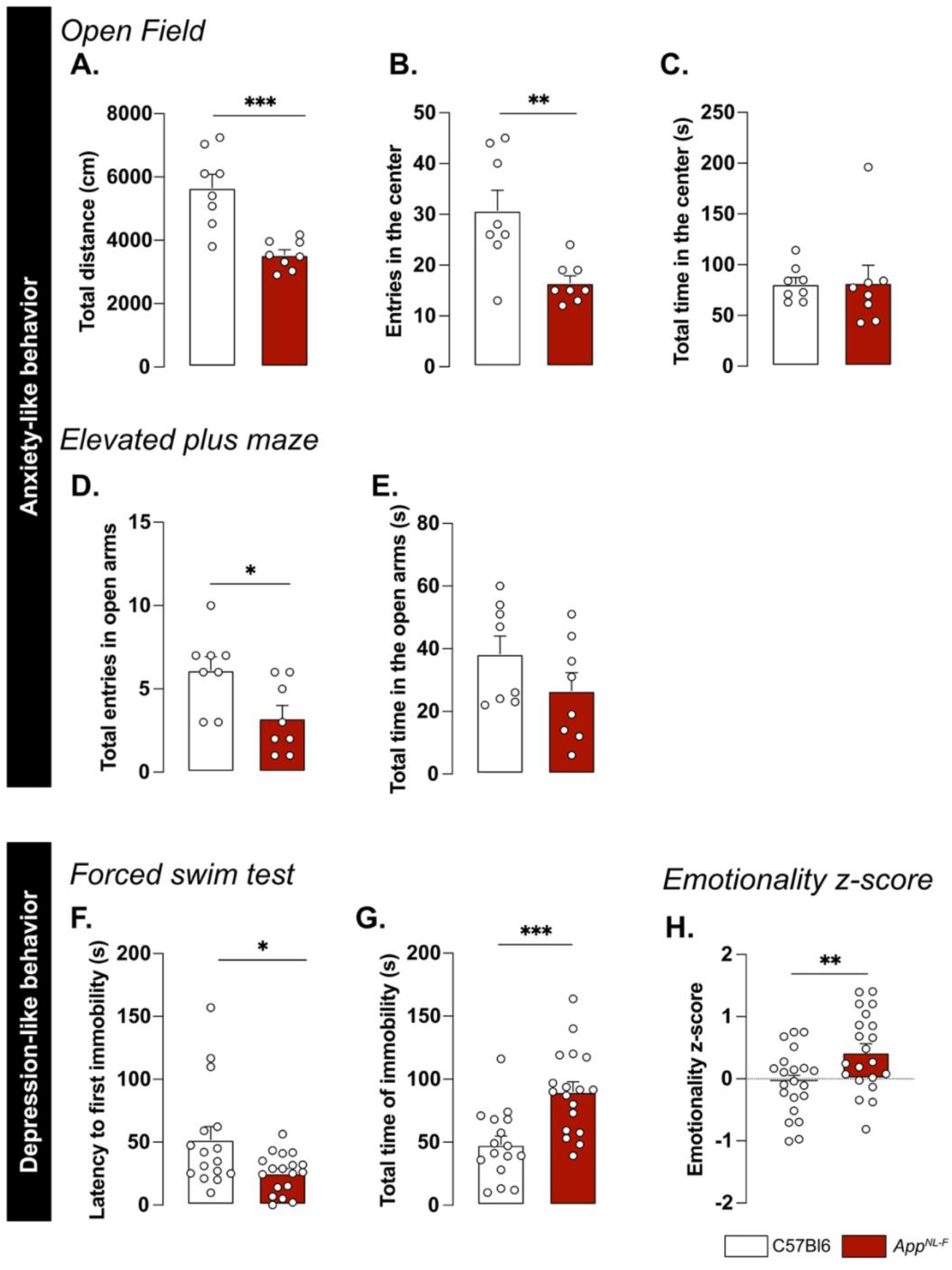
AppNL^-F^ mice display mild depressive-like behavior. In the open filed test, the total travelled distance (A.) is reduced in App^NL-F^ compared to C57Bl6, as well as the number of entries in the central area (B.). The total time in the open area is not different between the two strains (C.). Mann-Whitney test: **, p<0.01, ***, p<0.001 statistically different as shown. C57Bl6: n=8, App^NL-F^: n=8. In the elevated plus maze, the number of entries in the open arms is significantly lower in App^NL-F^ compared to C57Bl6 (D.) whereas the total time spent in the open arms (E.) remains unchanged between genotypes. Student t-test: *, p<0.05 statistically different as shown. C57Bl6: n=8, App^NL-F^: n=8. In the forced swim test, the latency to first immobility is lower in App^NL-F^ mice compared to C57Bl6 (F.), while the total time of immobility is higher (G.). Mann-Whitney and student t-test: *, p<0.05, ***, p<0.001 statistically different as shown. C57Bl6: n=16, App^NL-F^: n=18. The emotionality z-score is an integrative score allowing comparison of multiple factors, calculated on different scales and includes parameters from the different behavioral tests described above (see supplementary information). The emotionality z-score is significantly higher in AppNL-F mice compared to C57BL6 (H.). Student t-test: **, p<0.01 statistically significant as shown. C57Bl6: n=21, App^NL-F^: n=21.

We then explored the depression-like behavior in three different paradigms. In the forced swim test, the immobility was used as an indicator of helplessness. The latency to the first immobility was significantly reduced in *App^NL-F^* mice (C57Bl6: 52.05 ± 0.30 seconds, *App^NL-F^*: 25.43 ± 3.70 seconds, p<0.05; fig. 5F) whereas the total time of immobility was significantly increased (C57Bl6: 48.07 ± 6.89 seconds, *App^NL-F^*: 90.07 ± 7.79 seconds; p<0.001; fig. 5G). In the splash test (supplementary methods), the lack of grooming reflects a lack of self-care. No significant differences in the grooming behavior were observed in the *App^NL-F^*mice compared to C57Bl6 (fig. S4A-C). The sucrose preference test (supplementary methods), where a lack of preference for sugar compared to water is considered to reflect anhedonia, revealed no difference between the two strains (fig. S4D).

These behavioral data suggest a slight anxiety– and depression-like behavior and this was quantified as an emotionality z-score [40]. We used several parameters per test to establish a z-score for each test (ZSc_test; Table S1) and each ZSc_test was then combined into an emotionality z-score (fig. 5H). The C57Bl6 mice were used as the control group and have an average z-score of zero (–0.05 ± 0.11). The *App^NL-F^* mice had a significantly increased z-score (0.42 ± 0.13, p<0.01) confirming depressive-like behavior in the six months old *App^NL-F^* mice.

### A single dose of ketamine (5mg/kg) partially corrects depressive-like behavior in App^NL-F^ mice

A low dose of ketamine has antidepressant effects, and can also revert dysfunction in synaptic tuning [44]. Thus, an i.p injection of either ketamine (5mg/kg) or vehicle solution (NaCl 0.9%) was administered to six months old *App^NL-F^* mice and behavior was tested twenty-four hours after injection. In the five-trials social memory test, ketamine did not affect social memory as shown by a similar decrease of social interaction over four consecutive trials, in both treatment conditions (fig. 6A). As in our previous experiments, the social interaction measured in the fifth trial was not significantly different from the fourth trial in the vehicle treated animals (trial 4: 18.90 ± 4.66 % vs trial 5: 38.20 ± 5.91 %, fig. 6B). Ketamine did not have a significant effect on the social interaction on the fifth trial (trial 4: 22.085 ± 4.97 % vs. trial 5: 40.12 ± 9.48 %).

**Figure 6:**
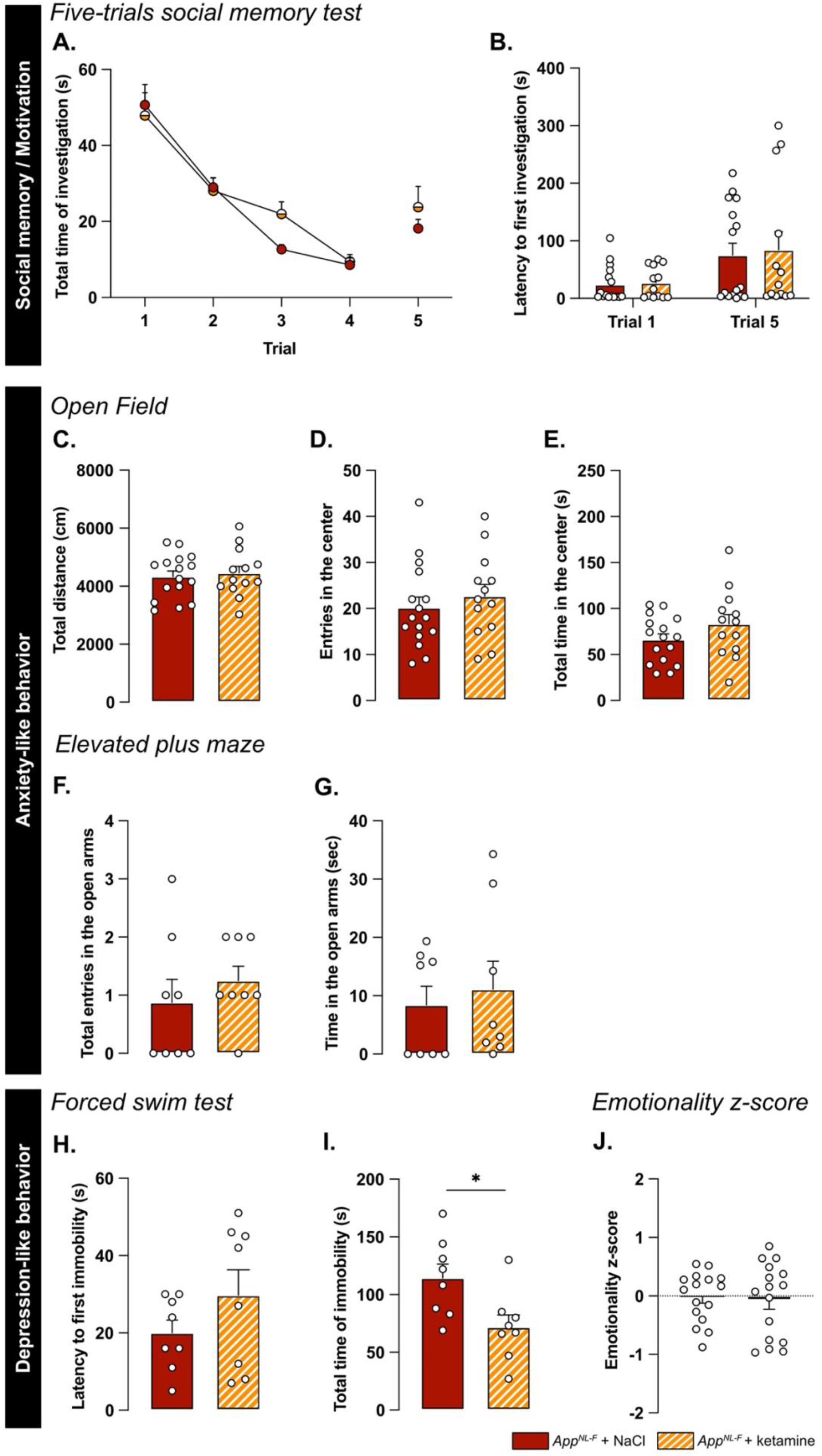
Ketamine (5mg/kg) partially restores emotionality in six-months old App^NL-F^ mice. As previously shown, App^NL-F^ mice do not have social memory impairments, as the social interaction progressively declines over four trials. This learning curve is not affected by injections of NaCl of Ketamine twenty-four hours before testing (A.). Moreover, ketamine does not significantly affect the lack of motivation (B.). App^NL-F^ + NaCl: n=16, App^NL-F^ + ketamine: n=13. In the open filed test, neither the total travelled distance (C.), the number of entries in the open area (D.) nor the total time in that zone (E.) is affected by ketamine. App^NL-F^ + NaCl: n=16, App^NL-F^ + ketamine: n=13. A similar profile was obtained in the elevated plus maze in which neither the number of entries in the open arms (F.) or the total time in that zone (G.) are affected. App^NL-F^ + NaCl: n=8, App^NL-F^ + ketamine: n=8. In the forced swim test, the latency to first immobility (H.) is unchanged after ketamine treatment while the total time of immobility (I.) is reduced. Student t-test: *, p<0.05 statistically different as shown. App^NL-F^ + NaCl: n=8, App^NL-F^ + ketamine: n=8. Interestingly, the emotionality z-score (J.) remains unchanged after ketamine 5mg/kg treatment. App^NL-F^ + NaCl: n=16, App^NL-F^ + ketamine: n=16.

We further explored the antidepressant action of a single injection of 5mg/kg ketamine. In the open field, none of the parameters total distance travelled (NaCl: 4331 ± 190.5 cm, ketamine: 4454 ± 229.3 cm; fig. 6C), the number of entries in the center (NaCl: 20.19 ± 2.30 entries, ketamine: 22.69 ± 2.55 entries; fig. 6D) and total time spent in that zone (NaCl: 66.11 ± 6.35 seconds, ketamine: 83.13 ± 10.32 seconds; fig. 6E) were changed by the ketamine treatment. Likewise, ketamine had no effect on the number of entries in the open arms in the elevated plus maze (NaCl: 0.87 ± 0.40 entries, ketamine: 1.25 ± 0.25 entries; fig. 6F) nor the total time spent in that zone (NaCl: 8.41 ± 3.2 seconds, ketamine: 1.12 ± 4.79 seconds; fig. 6G).

In the forced swim test, no changes were observed in the latency to first immobility (NaCl: 20 ± 3.32 seconds, ketamine: 29.75 ± 6.56 seconds; fig. 6H). However, there was a reduced total time of immobility in ketamine treated mice compared to NaCl treated (NaCl: 114.4 ± 2.02 seconds, ketamine: 71.88 ± 10.61, p<0.05; fig. 6I). Similarly, in the splash test, a reduced latency to first grooming (NaCl: 32.52 ± 3.71 seconds, ketamine: 16.30 ± 2.56 seconds, p<0.01; fig. S4E), increased frequency of grooming (NaCl: 7 ± 0.27, ketamine: 9.13 ± 0.69, p<0.05; fig. S4F) and an increased total time of grooming (NaCl: 145.7 ± 6.01 seconds, ketamine: 188.8 ± 3.56 seconds, p<0.05; fig. S4G) were observed after ketamine treatment.

The behavioral data was integrated into the emotional z-score (fig. 6J) to assess the overall effect of ketamine on the depressive-like behavior of *App^NL-F^*. Animals who received the vehicle solution were used as the control group with an average emotionality z-score of zero. No statistical difference was shown in the z-score between the two different groups. As expected, ketamine had an antidepressant effect in *App^NL-F^* mice. This effect, however, did not target the specific behaviors that are affected in *App^NL-F^*mice, but was a general effect.

### A single dose of ketamine (5mg/kg) partially restores synaptic dysfunctions and reduces astrocytic tonic inhibition

Twenty-four hours after either ketamine or vehicle solution, no difference was observed on resting membrane potential (NaCl: –53.4 ± 3.08 mV, ketamine: –51.8 ± 5.29 mV; fig. 7A) or any sEPSC parameters (fig. S3M-P). However, ketamine induced a significant increase in frequency of mEPSC (NaCl: 0.10 ± 0.001 Hz, ketamine: 0.59 ± 0.11 Hz, p<0.01; fig. 7B) while the amplitude (NaCl: –29.03 ± 4.40 pA, ketamine: –21.89 ± 3.52 pA; fig. 7C), decay time (fig. S3Q) and rise time (fig. S3R) were not affected.

**Figure 7:**
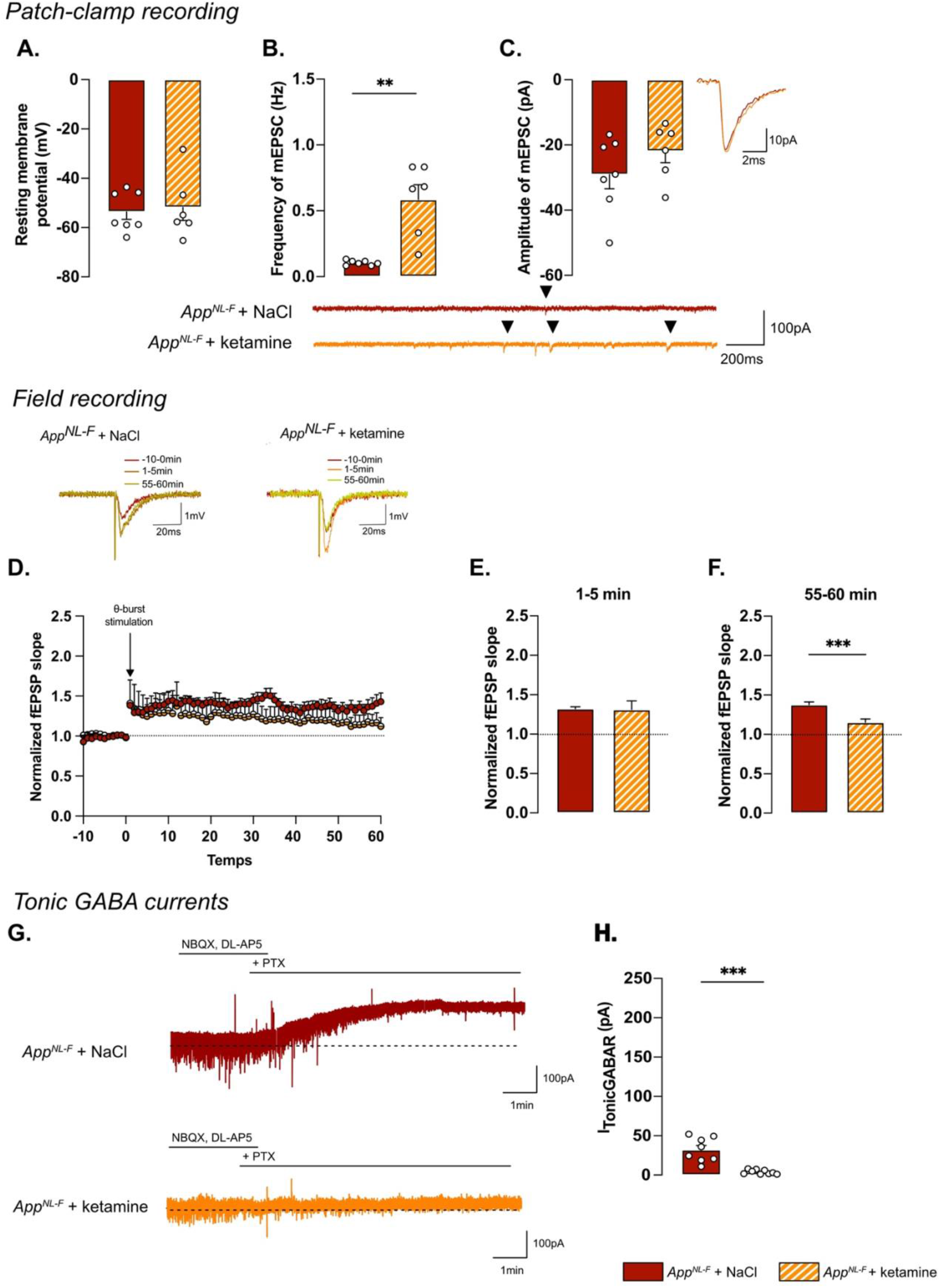
Synaptic mistuning in App^NL-F^ is restored by a single dose of ketamine (5mg/kg) A single injection of ketamine twenty-four hours before recordings did not affect resting membrane potential in App^NL-F^ mice (A.). In the same cell, the frequency of mEPSC is increased by ketamine treatment (B.)., whereas the amplitude of mEPSC remains unchanged (C.). Mann-Whitney test: **, p<0.01 statistically different as shown. App^NL-F^ + NaCl: n=7 cells recorded in five animals, App^NL-F^ + ketamine: n=6 cells recorded in six animals. With the same θ-burst stimulation as previously used, the fEPSP magnitude goes back nearly to the baseline value in App^NL-F^ + ketamine mice whereas in the App^NL-F^ + NaCl, the stimulation triggers a long-term potentiation (D.). Representative traces are shown on the top. There is no difference between in the fEPSP magnitude in the two groups at 0-5 minutes after stimulation (E.). However, at 55-60 minutes after the stimulation the fEPSP magnitude is significantly higher in NaCl treated mice compared to ketamine treated mice (F.) Student t-test: ***, p<0.001 statistically different as shown. App^NL-F^ + NaCl: n=5 recordings in four animals, App^NL-F^ + ketamine: n=5 recordings in four animals. Tonic GABA currents were recorded (G) and the average current was significantly reduced in App^NL-F^ mice treated with ketamine compared to NaCl treated mice (H.). Mann-Whitney test ***, p<0.001 significantly different as shown. App^NL-F^ + NaCl: n=8 cells recorded in four animals; App^NL-F^ + ketamine: n=9 cells recorded in four animals.

Synaptic plasticity deficits in six months old *App^NL-F^*mice were restored by ketamine: a subthreshold θ-burst stimulation gave an initial potentiation of the fEPSP in both conditions (NaCl: 1.32 ± 0.02, ketamine: 1.31 ± .11; fig. 7D-E). Fifty-five minutes after stimulation, the magnitude of the long-term potentiation was significantly smaller in *App^NL-F^* + ketamine compared to *App^NL-F^* + NaCl (fig. 7D-F; p<0.001). Taken together, these electrophysiological recordings confirm the ability of ketamine to retune misregulated synaptic transmission and plasticity.

Since we hypothesize that the synaptic mistuning in *App^NL-F^* mice is mediated through astrocytes, we tested whether ketamine would affect the astrocytic tonic inhibition (fig. 7G). Tonic GABA currents were recorded in hippocampal slices from *App^NL-F^* mice that had been treated with ketamine twenty-four hours earlier. Tonic GABA currents were significantly decreased in ketamine treated *App^NL-F^* mice compared to saline treated group (3.94 ± 0.9 pA vs. 32.11 ± 5.457 pA, p<0.001; fig. 7H), suggesting a role for astrocytes in the antidepressant effect of ketamine.

## DISCUSSION

As the final product of the dysregulated amyloid cascade, Aβ plaques have traditionally been studied as one of the main cause of Alzheimer’s pathology [48], although we know that both behavioral and pathophysiologal changes occur before plaques can be detected [49]. In a late onset mouse model of Alzheimer’s disease (*App^NL-F^*), at six months of age, Aβ plaques are not detectable, but there is an increase in non-soluble dimers. Here we show that at this age, *App^NL-F^* mice have reactive-like astrocytes, increased tonic GABA, and synaptic misfunction. This is in good agreement with the hypothesis that Aβ oligomers are inducing several toxic events in early Alzheimer’s disease [50].

A reduced threshold for long-term potentiation in *App^NL-F^*mice is a surprising finding. Memory impairments and dementia disorders have typically been associated with a decrease in synaptic potentiation; previous work in the *App^NL-F^* mice at six months of age show no effect on LTP after a strong tetanic stimulation [51]. However, synaptic potentiation is not linearly correlated with memory, rather synaptic strength is constantly adjusted based on activity patterns in the neuronal network [52]. In order to learn and adapt to the changing environment, synaptic transmission needs to be plastic, while keeping the overall activity within optimal range. In fact, it has been proposed that cognitive deficits in early Alzheimer’s disease is not about reduced plasticity, but an imbalance at the network level [53], which is compatible with a reduced threshold for potentiation. Our results in the *App^NL-F^* mice are consistent with previous work, where authors have been using a low or even sub-threshold stimulation protocol in other model with memory impairment [20]. The increase in synaptic potentiation, as well as the lack of change in evoked synaptic response, in six months old *App^NL-F^* mice has also been described by others [19], who attribute the effect to a decreased turnover of presynaptic proteins.

In this work, we reveal that blocking GABA synthesis with the MAO-B blocker deprenyl changes synaptic transmission in *App^NL-F^*mice in several ways. As expected, deprenyl pre-treatment increases the amplitude of synaptic events, both when evoked by spontaneous action potentials (sEPSC) and when arising from non-evoked release of individual vesicles (mEPSC). This effect is easily explained by an increase in membrane resistance due to the decreased GABA tone. In agreement, the frequency of mEPSC is increased in deprenyl-treated slices, however the frequency of sEPSC is not. Interestingly, the threshold for LTP is increased by blocking MAO-B and not reduced as one would expect with a reduced GABA tone. Thus, blocking GABA synthesis in astrocytes in *App^NL-F^*mice entails more and other changes than a direct increase in membrane resistance. Previous work in our lab has shown that synaptic activity and plasticity are strongly interacting in a well-tuned relationship [44]. Thus, the increased synaptic activity induced by deprenyl could tune synaptic plasticity to specifically reverse synaptic deficits in the *App^NL-F^* mice.

The synthesis and release of GABA from reactive astrocytes are well described phenomena and were first shown in a mouse model overexpressing App and with a severe plaque load [54]. Here we describe that astrocyte morphology is changed and tonic GABA is increased already before plaque formation in the *App^NL-F^* mice model of Alzheimer’s disease. Reactive astrocytes [28] have been described as the starting point of several brain dysfunctions and proposed as an interesting therapeutic target in various brain disorders [55], and more recently in neurodegeneration and aging [56]. Although we do not directly rule out a neuronal origin of GABA, the astrocytic origin of GABA is strongly supported by results shown that the main effect of blocking MAO-B in the hippocampus is a reduction in astrocytic GABA [42, 57].

Reactive astrocytes, as well as synaptic impairments, are linked to depressive-like behavior [58]. *App^NL-F^* mice do indeed display depressive like behaviors in some tests already at six months. There is no test that directly translates to clinical depression, partly due to the fact that clinical depression can be describes as a syndrome, with different clinical presentations [59]. To get an overall vision of the emotional state of the *App^NL-F^* mice, we compiled our behavioral data into an emotionality z-score, relevant to study depression-like behaviors assessed on different scales [60]. Since the reduced mobility could be another confounding factor in several of the tests used to assess depression-like behavior, data obtained in the open field and elevated plus maze were normalized with the travelled distance. When all the affective behavioral test are integrated in a z-score there is a significant difference in *App^NL-F^* compared to C57Bl6 mice. This is in good agreement with early depressive symptoms observed in Alzheimer’s patients [10] as well as results from other animals models of the disease [61]. Moreover, we confirm the lack of memory impairment at this age using the five-trials social memory test. In the last session of this test, the *App^NL-F^*do not show increased time of interaction with the novel individual, as does the C57Bl6 mice. Having ruled out that the lack of interest is due to olfactory problems, we suggest that this lack of interaction is due to an apathy-like behavior, which is consistent with the depressive like phenotype. However, another explanation could be that *App^NL-F^*mice have deficits in social recognition consistent with recent observation in early diagnose patients [62]. We here use as the “presenting animal”, mice with the same fur color, same age, same gender as the tested animal. Thus, *App^NL-F^*mice might present early pattern separation deficits, as they cannot discriminate two very close situations (i.e., identify the mice as a new individual), in line with deficits observed in early Alzheimer’s disease [63]. Further work will need to be done to clarify this.

Finally, we investigated the therapeutic effect of a low dose of ketamine (5mg/kg) delivered i.p, twenty-four hours prior to experiments. In line with the literature [64], we successfully show the antidepressant effect of this drug in the forced swim test and the splash test. There was no significant effect in the anxiety related tests and the overall emotionality z-score was not significantly improved. Moreover, the lack of exploration of the novel mice at the fifth trial in the social exploration test was not affected by ketamine. The partial effect of ketamine on behavior is in contrast to the significant effect of ketamine on the synaptic dysfunction in the *App^NL-F^* mice, where mEPSC frequency is increased and the threshold for LTP induction is restored. This is in good agreement with previous work from our lab where we show that ketamine affects synaptic transmission and plasticity in a state-dependent manner, to stabilize the synaptic tuning [44]. Thus, a single dose of ketamine corrects early synaptic dysfunctions, in the *App^NL-F^* mice, but do not specifically reverse behavioral deficits. However, ketamine does have a general antidepressant effect on several of the behaviors tested. It is interesting to note that deprenyl, that also reverts synaptic dysfunctions in the *App^NL-F^* mice, has been used clinically as an anti-depressant drug [65], although this effect has typically been ascribed to its effect on monoamine synthesis rather than the effect on astrocytes.

Increasing synchronous neuronal activity in the gamma frequency has been shown to slow-down disease progress and improve cognitive performance in Alzheimer’s disease [66] (but see also [67]). Moreover, recent work suggest that neuronal activity can indeed affect Aβ accumulation and plaque deposition [68]. Restoring synaptic transmission and plasticity may thus have a prophylactic effect on Alzheimer disease pathology. Furthermore, we can speculate that restoring synaptic tuning may be a biological mechanism underlying the beneficial effect of a mental and physical active life-style [69]. Thus, it would be interesting to follow-up the long-term effect of chronic ketamine treatment, with its beneficial effect on synaptic tuning on the progression of Alzheimer’s disease.

Ketamine’s mechanism of action is still debated. Interestingly, we see that ketamine and deprenyl have the same effects on synaptic transmission and plasticity in *App^NL-F^* mice. Furthermore, we show that ketamine reduces tonic GABA, thus one possibility is that ketamine acts on reactive astrocytes to restabilize synaptic tuning, through changes in gliotransmission. This is in good agreement with recent results showing that ketamine affects astrocytic morphology and function [70], as well as astrocytic GABA metabolism [71].

Research on Alzheimer’s disease treatment has focused mainly on Aβ reducing treatments [72] or the cholinergic system [73]. Here we show that targeting the glutamatergic system can correct early synaptic impairment as well as early depression-like behaviors observed in an Alzheimer’s disease model. How an early intervention restoring synaptic function affects the progression of Alzheimer’s disease remains to be shown.

## AUTHOR CONTRIBUTION

Conceptualization Maria Lindskog and Per Nilsson; Investigation Benjamin Portal, Moa Södergren, Teo Parés I Borrell, Romain Giraud and Nicole Metzendorf; Supervision Greta Hultqvist and Maria Lindskog; Writing the original draft preparation Benjamin Portal and Maria Lindskog; Funding acquisition Benjamin Portal and Maria Lindskog.

## INSTITUTIONAL REVIEW BOARD STATEMENT

The study was conduct in accordance with the Declaration of Helsinki and was approved by the Swedish board of animal use (Jordbruksverket ethical permit #5.2.18-03389/2020) and accepted by the ethics committee at Uppsala University (Djurskyddsorganet at Uppsala University, accreditation #DOUU-2020-022).

## Supporting information

Portal et al. – Supplementary information

## LIST OF ABBREVIATIONS

Aβ: Amyloid-beta
aCSF: Artificial cerebrospinal fluid
App: Amyloid precursor protein
DL-AP5: DL (2*R*)-amino-5-phosphonovaleric acid
fEPSP: Field excitatory post-synaptic potential
GABA: Gama aminobutyric acid
GFAP: Glial fibrillary acidic protein
i.p.: Intraperitoneal
LTP: Long term potentiation
MAO-B: Monoamine oxidase type B
mEPSC: Mini excitatory post-synaptic current
NaCl: Natrium chloride
NBQX: 2,3-dioxo-6-nitro-7-sulfamoyl-benzo[f]quinoxaline
PTX: Picrotoxin
sEPSC: Spontaneous excitatory post-synaptic current
TTX: Tetrodotoxin

## ACKNOWLEDGMENTS

The authors would like to thank both the BioVis imaging facility and the Uppsala University Behavioral Facility (UUBF) for their expertise and their help on setting up the different experiments. The authors would like to thank Takaomi Saido and Takashi Saito at RIKEN Center for Brain Science (Japan) for providing the *App^NL-F^*mice.

