## Supplementary material for "Early astrocytic dysfunction is associated to mistuned synapses as well as anxiety and depressive-like behavior in the *App^NL-F^* mouse model of Alzheimer’s disease": Portal et al. – Supplementary information

### SUPPLEMENTARY METHODS

*Amyloid-beta (Aβ) ELISA.*

*Perfusion and brain extraction.*

Mice were anesthetized with a mixture of ketamine (100 mg/kg) and xylazine (16 mg/kg) administered via i.p route. In absence of corneal reflexes, mice were transcardially perfused with NaCl 0.9%. The brain was extracted, and hippocampi, cortex, and cerebellum were dissected in dry ice and kept at -80°C until further use.

Brain homogenization.

The brain regions were homogenized at 1:5 weight/volume ratio in precellys homogenization tubes (Bertin Technologies, P000933-LYSK0-A). A three-step homogenization was applied: Tris-buffered saline (TBS) to isolate soluble Aβ, TBS with 1% triton-x100 (TBS-T; Sigma-Aldrich, X100) for isolation of membrane-associated Aβ, and 70% (v/v) formic acid (FA; Sigma-Aldrich, F0507) for the insoluble Aβ fraction. TBS-homogenized brains were centrifuged at 16 000 x g for sixty minutes at +4°C, the supernatant was collected. The pellets were subsequently homogenized in TBS-T and centrifuged at 16 000 x g for sixty minutes at +4°C and supernatant was collected. The pellets were dissolved in 70% (v/v) FA, centrifuged at 16 000 x g for sixty minutes at +4°C and the supernatant was collected and immediately neutralized with 1M Tris pH 8.0 (Thermo Fisher Scientific, AM9856).

Aβ ELISA.

The three Aβ fractions were analyzed using a homogenous sandwich ELISA using the Aβ N-terminal specific antibody 3D6 (murine version of Bapineuzumab, produced in-house) as both capture and detection antibody. 96-wells ELISA plates were coated with 1 µg/mL 3D6 and incubated over night at +4˚C. Plates were blocked with 1% (w/v) BSA in PBS 1X for two hours. Dilutions of TBS and TBS-T brain fractions (1:10 (v/v)) and FA brain fraction (1:200 (v/v)) in ELISA incubation buffer (EIB; 0.05% tween-20 (Sigma-Aldrich, P9416), 0.1% w/v BSA in PBS 1X) were added to the ELISA plate and incubated over nigh at +4˚C. Biotinylated 3D6 (prepared in-house; 1 µg/mL in EIB) was incubated for two hours at room temperature followed by one hour of streptavidin-HRP (Mabtech, 3310-9-1000) at room temperature. The ELISA was developed with K-blue Aqueous TMB substrate (Neogen Corp., 331177) and read with a spectrophotometer (Tecan, Spark) at 450 nm.

*Behavioral analysis*

Splash test.

The animals were isolated in a new cage and habituated for at least twenty minutes. The test was performed as previously described (Quesseveur et al., 2015): 200 µL of 10%

sucrose solution was squirted on the mouse's snout and the grooming behavior (latency to first, frequency and total time), used to evaluate self-care, was manually scored for five minutes. Mice were placed back together in their home cage after the experiment.

Sucrose preference test.

Each animal was singled housed and habituated to drink water from two identical bottles for forty-eight hours. During the following seventy-two hours, mice could choose between water and a 1% sucrose solution. Sucrose solution intake was measured by weighing the bottles before and after the last twenty-four hours and expressed as the percentage of the total amount of liquid ingested, normalized to the mouse body weight. Each bottle was replaced and switched from left to right daily.

### SUPPLEMENTARY FIGURES

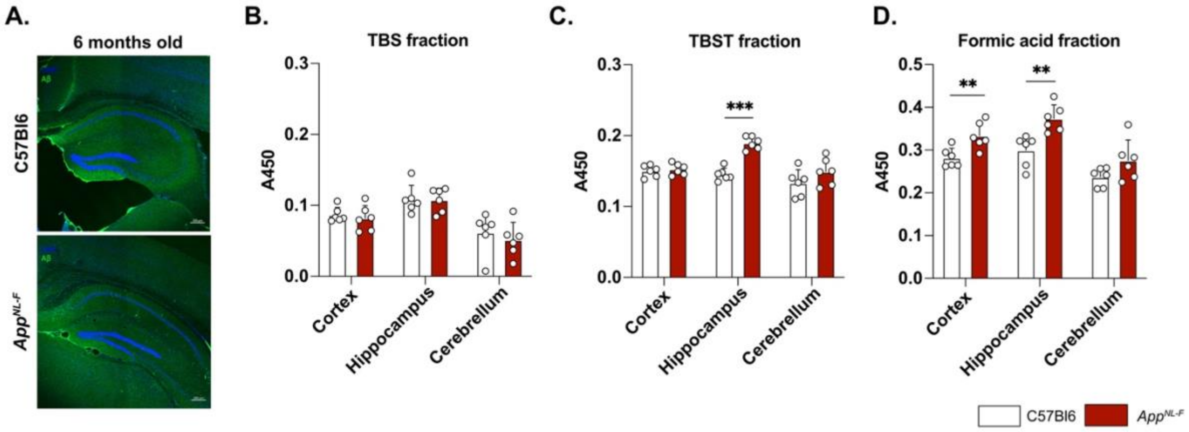


**Figure S1: *$App^{NL-F}$ mice do not display Aβ plaques but higher levels of Aβ in the insoluble fraction in the hippocampus.***

*Immunolabelling of Aβ (A.) reveals the absence of plaques in the hippocampus of six months old $App^{NL-F}$ mice, compared to C57Bl6 mice. ELISA analysis of different Aβ fractions shows unchanged concentrations in non-membrane-bound Aβ (TBS fraction, B.) in the hippocampus, the cortex and the cerebellum. However, increased concentration of membrane-bound Aβ (TBST fraction) in the hippocampus (C) but not in the cortex nor the cerebellum were observed. Our analysis also revealed an increased formic acid fraction (i.e., non-soluble dimers) in both the cortex and the hippocampus (D.).*

*For the three fractions: Two-way ANOVA with the genotype and the brain region as the main factors. Bonferroni multiple comparison: **, $p<0.01$; ***, $p<0.001$ statistically significant as shown. C57Bl6: n=6, $App^{NL-F}$: n=6. Figure A: scale bar 200µm.*

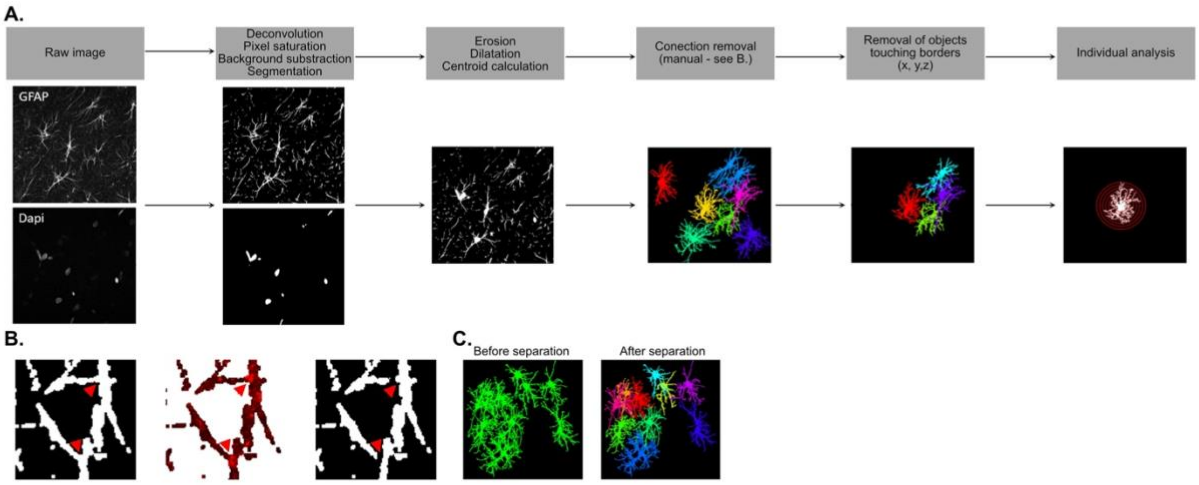


***Figure S2: Astrocyte morphology analysis workflow*** *(related to fig. 1)*
*Z-stack images of GFAP and DAPI stainings were merged in a z-projection and were treated according the presented workflow (A.). Before reconstruction astrocytes were manually separated (B.): the first image shows the connection (red arrows) on the deconvoluted image while the second image shows the raw image of the connexin 43 (Cx43) staining, use to discriminate connecting hubs between astrocytes. The third image shows the absence of connections after manual removal. (C.) depicts how the syncytium looks like before, and after separation. Once all astrocytes have been separated, the morphology of each cell was 3D reconstructed and individually analyzed.*

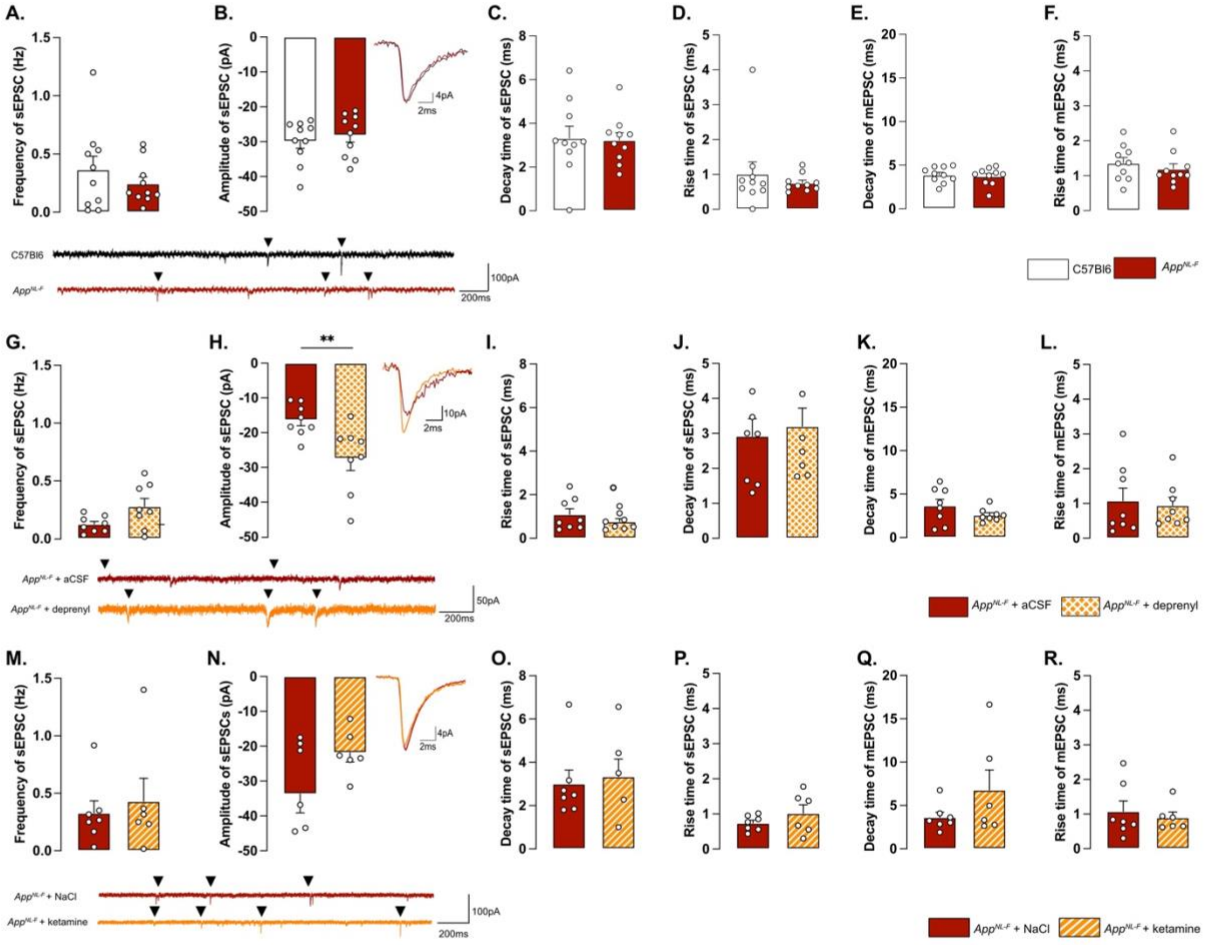


***Figure S3 : Complementary electrophysiological measurements (related to figure 2, 3 and 7)***

*Patch clamp experiments revealed no difference in frequency (A.), amplitude (B.) decay time C.) or rise time (D.) of sEPSC. No differences were unveiled either in decay time (E.) nor rise time (F.) of mEPSC. Representative traces for sEPSC are depicted under the graphs. C57Bl6 controls: n=10 cells recorded in seven animals, App$^{NL-F}$: n=10 cells recorded in seven animals. Pre-treatment with deprenyl didn't affect the frequency of sEPSC (G.) but increased the amplitude (H.) Representative traces are depicted under the graphs. Neither the rise time or the decay time for sEPSC (I. and J. respectively) and mEPSC (K. and L. respectively) were affected by deprenyl. App$^{NL-F}$ + aCSF: n=8 cells recorded in four animals; App$^{NL-F}$ + deprenyl: n=8 cells recorded in four animals.*

*No differences were unveiled in frequency (M.), amplitude (N.), decay time (O.) or rise time (P.) of sEPSC after ketamine treatment. The NMDA receptor blocker didn't affect the decay time (Q.) nor the rise time (R.) of mEPSC. Representative traces for sEPSC are depicted under the graphs. App$^{NL-F}$ + NaCl: n=7 cells recorded in five animals, App$^{NL-F}$ + ketamine: n=6 cells recorded in six animals.*

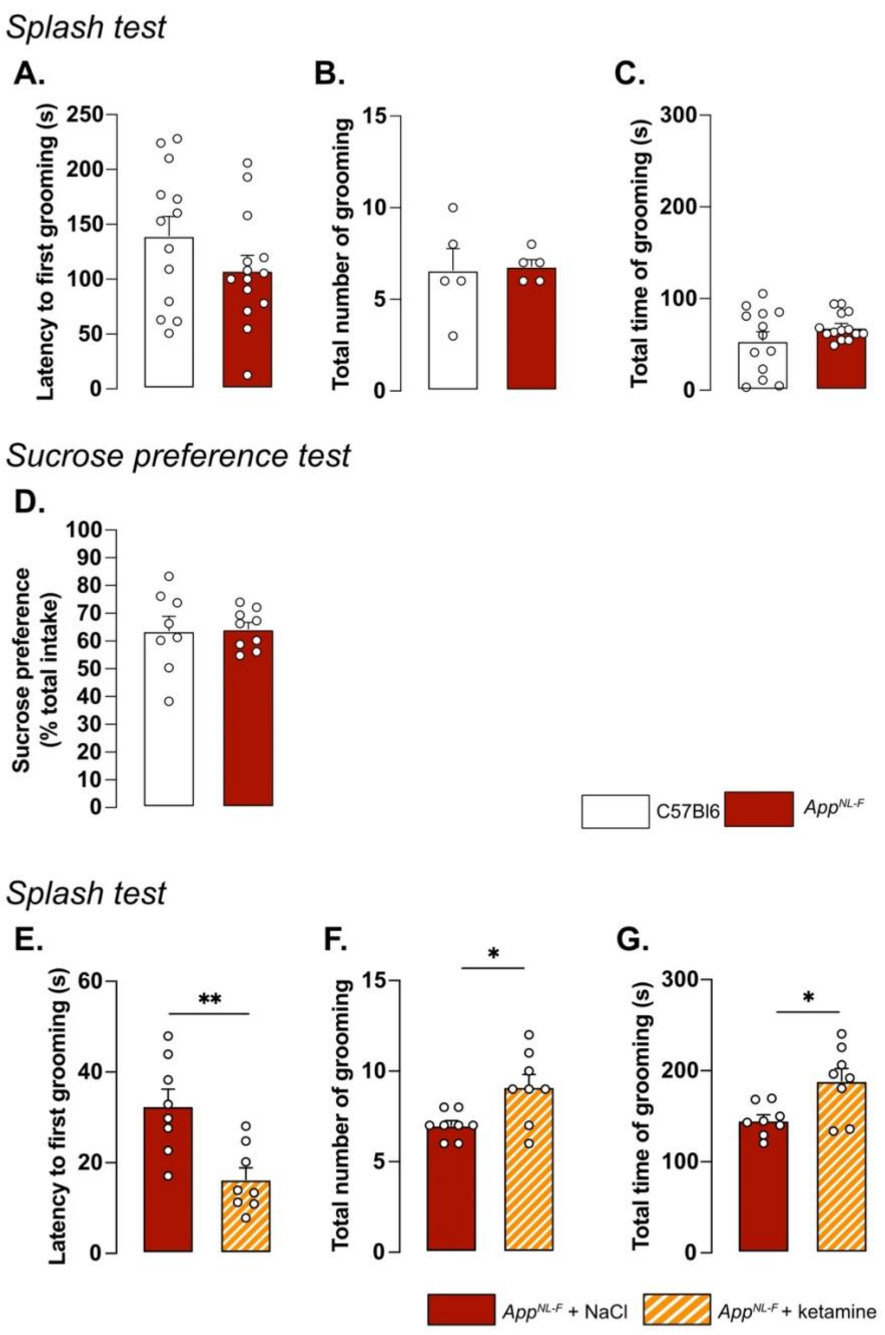


**Figure S4 : *Splash test and sucrose preference test*** *(related to figure 5 and 6)*

*No differences were unveiled in the latency to first grooming (A.), total number of grooming (B.) nor in the total time of grooming (C.). C57Bl6: n=13, App$^{NL-F}$: n=14. For the frequency n=4 and n=5 for C57Bl6 and App$^{NL-F}$ respectively. In the sucrose preference test, no difference was unveiled in the sucrose consumption (D.) C57Bl6: n=8, App$^{NL-F}$: n=9.*

*After a single injection of ketamine (5mg/kg), the latency to first grooming (E.), total number of grooming (F.) and in the total time of grooming (G.) were increased, compared to saline treated animals. Student t-test: *, p<0.05, **, p<0.01 statistically different as shown. App$^{NL-F}$ + NaCl: n=8, App$^{NL-F}$ + ketamine: n=8.*

127

*Table S1*

| | | zSc_param | | | | | | | | | | | | | zSc_test | | | | | |
|---|---|---|---|---|---|---|---|---|---|---|---|---|---|---|---|---|---|---|---|---|
| Cohort | Genotype | zSC_OF_Distance | zSC_OF_Center entries | zSC_OF_Center time | zSC_OF_Center/Total distance ratio | zSC_EPM_Open arms entries | zSC_EPM_Open arms time | zSC_EPM_Open/Total arms entries | zSC_ST__Grooming latency | zSC_ST_Grooming frequency | zSC_ST_Grooming duration | zSC_TST_Immobility latency | zSC_TST_Immobility duration | zSC_SPT_Sucrose consumption | zSC_OF | zSC_EPM | zSC_ST | zSC_TST | zSC_SPT | Emotionality z-score |
| *Formula* | | *(X-μ)/σ* | | | | | | | | | | | | | *Average parameter (per test)* | | | | | *Average z-score* |
| 1 | C57Bl6 | 0,49 | 0,42 | -1,88 | 1,59 | 0,50 | -0,26 | 0,57 | | | | -0,64 | -1,73 | | 0,15 | 0,27 | | -1,19 | | -0,25 |
| 1 | C57Bl6 | 0,95 | 0,60 | -0,85 | 1,32 | -2,18 | -1,55 | -1,53 | | | | 1,84 | 0,17 | | 0,51 | -1,75 | | 1,00 | | -0,08 |
| 1 | C57Bl6 | 1,56 | 1,59 | 1,03 | 0,34 | -0,17 | -0,79 | -0,47 | | | | -0,20 | -0,48 | | 1,13 | -0,48 | | -0,34 | | 0,10 |
| 1 | C57Bl6 | -1,16 | -0,83 | 0,58 | -1,03 | -0,17 | -0,26 | -0,64 | | | | -1,00 | 0,27 | | -0,61 | -0,35 | | -0,37 | | -0,44 |
| 1 | C57Bl6 | 0,22 | 0,42 | 0,48 | -0,31 | 1,18 | 1,57 | 1,35 | | | | -0,49 | -0,55 | | 0,20 | 1,37 | | -0,52 | | 0,35 |
| 1 | C57Bl6 | -0,37 | 0,25 | 1,02 | -0,97 | -0,17 | -0,26 | -0,84 | | | | 0,82 | 0,37 | | -0,02 | -0,42 | | 0,59 | | 0,05 |
| 1 | C57Bl6 | -1,33 | -1,27 | -0,21 | -0,68 | 0,50 | 0,50 | 0,77 | | | | 0,60 | 1,79 | | -0,87 | 0,59 | | 1,20 | | 0,31 |
| 1 | C57Bl6 | -0,37 | -1,18 | -0,16 | -0,25 | 0,50 | 1,03 | 0,77 | | | | -0,93 | 0,17 | | -0,49 | 0,77 | | -0,38 | | -0,03 |
| 2 | C57Bl6 | | | | | | | | -0,24 | | -0,80 | -0,16 | 0,40 | 0,17 | | | -0,52 | 0,12 | 0,17 | -0,08 |
| 2 | C57Bl6 | | | | | | | | 1,05 | | -0,49 | -0,69 | 0,38 | 0,85 | | | 0,28 | -0,15 | 0,85 | 0,33 |
| 2 | C57Bl6 | | | | | | | | 0,74 | | 1,09 | -0,87 | 0,53 | -1,74 | | | 0,92 | -0,17 | -1,74 | -0,33 |
| 2 | C57Bl6 | | | | | | | | 0,17 | | 0,48 | 0,76 | -0,32 | 1,34 | | | 0,33 | 0,22 | 1,34 | 0,63 |
| 2 | C57Bl6 | | | | | | | | -1,01 | | -1,00 | -1,17 | 1,57 | -0,91 | | | -1,01 | 0,20 | -0,91 | -0,57 |
| 2 | C57Bl6 | | | | | | | | 0,10 | | -0,97 | 0,89 | -1,27 | -0,24 | | | -0,43 | -0,19 | -0,24 | -0,29 |
| 2 | C57Bl6 | | | | | | | | 0,99 | | 0,03 | 1,67 | -1,48 | -0,17 | | | 0,51 | 0,09 | -0,17 | 0,14 |
| 2 | C57Bl6 | | | | | | | | -1,80 | | 1,66 | -0,44 | 0,19 | 0,69 | | | -0,07 | -0,13 | 0,69 | 0,16 |
| 3 | C57Bl6 | | | | | | | | -0,98 | 1,38 | -1,63 | | | | | | -0,41 | | | -0,41 |
| 3 | C57Bl6 | | | | | | | | -1,26 | -1,30 | 0,35 | | | | | | -0,74 | | | -0,74 |
| 3 | C57Bl6 | | | | | | | | -0,48 | 0,23 | 0,57 | | | | | | 0,11 | | | 0,11 |
| 3 | C57Bl6 | | | | | | | | 0,05 | 0,23 | -0,21 | | | | | | 0,02 | | | 0,02 |
| 3 | C57Bl6 | | | | | | | | -1,56 | -0,54 | 0,92 | | | | | | -0,39 | | | -0,39 |
| 1 | *App$^{NL-F}$* | 1,79 | 1,67 | 2,18 | -0,60 | 1,18 | 1,57 | 1,35 | | | | -2,32 | -0,34 | | 1,26 | 1,37 | | -1,33 | | 0,43 |
| 1 | *App$^{NL-F}$* | 1,43 | 0,60 | 0,63 | 0,57 | -0,17 | -0,26 | -0,21 | | | | -2,10 | 1,89 | | 0,81 | -0,21 | | -0,10 | | 0,16 |
| 1 | *App$^{NL-F}$* | 1,25 | 1,05 | 1,13 | -0,06 | -0,17 | 0,28 | 0,06 | | | | 0,53 | 0,33 | | 0,84 | 0,06 | | 0,43 | | 0,44 |
| 1 | *App$^{NL-F}$* | 2,32 | 1,59 | 2,08 | 0,07 | 0,50 | 0,28 | 0,14 | | | | -0,57 | 0,91 | | 1,51 | 0,31 | | 0,17 | | 0,66 |
| 1 | *App$^{NL-F}$* | 1,97 | 1,41 | -6,53 | 9,29 | -0,84 | -1,76 | -0,99 | | | | -1,37 | 1,93 | | 1,53 | -1,20 | | 0,28 | | 0,21 |
| 1 | *App$^{NL-F}$* | 1,45 | 1,05 | -0,16 | 1,34 | -0,17 | -0,04 | -1,08 | | | | 0,09 | 0,57 | | 0,92 | -0,43 | | 0,33 | | 0,27 |
| 1 | *App$^{NL-F}$* | 1,83 | 1,41 | 0,21 | 1,53 | 0,50 | 0,93 | -0,21 | | | | -2,46 | 2,60 | | 1,24 | 0,41 | | 0,07 | | 0,57 |
| 1 | *App$^{NL-F}$* | 2,21 | 1,41 | -0,06 | 2,55 | 1,18 | 1,57 | 1,35 | | | | -0,64 | 0,81 | | 1,53 | 1,37 | | 0,09 | | 0,99 |
| 2 | *App$^{NL-F}$* | | | | | | | | -1,02 | | 0,56 | -0,89 | 4,11 | 0,24 | | | -0,23 | 1,61 | 0,24 | 0,54 |
| 2 | *App$^{NL-F}$* | | | | | | | | -0,16 | | 0,19 | -0,55 | 1,70 | 0,70 | | | 0,02 | 0,57 | 0,70 | 0,43 |
| 2 | *App$^{NL-F}$* | | | | | | | | 0,67 | | 0,69 | -1,23 | 8,42 | 0,58 | | | 0,68 | 3,60 | 0,58 | 1,62 |
| 2 | *App$^{NL-F}$* | | | | | | | | -1,16 | | 0,52 | -0,75 | 0,40 | -0,34 | | | -0,32 | -0,17 | -0,34 | -0,28 |
| 2 | *App$^{NL-F}$* | | | | | | | | -1,67 | | 0,62 | -0,52 | 0,98 | -0,61 | | | -0,52 | 0,23 | -0,61 | -0,30 |

| | | | | | | | | | | | | | | | |
|---|---|---|---|---|---|---|---|---|---|---|---|---|---|---|---|
| 2 | *App*$^{NL-F}$ | | | | 0,45 | | 0,34 | -1,08 | 5,44 | 0,17 | | 0,39 | 2,18 | 0,17 | 0,92 |
| 2 | *App*$^{NL-F}$ | | | | -1,54 | | 1,37 | -0,74 | 1,58 | -0,24 | | -0,08 | 0,42 | -0,24 | 0,03 |
| 2 | *App*$^{NL-F}$ | | | | -0,89 | | 1,37 | -0,27 | 3,91 | 0,39 | | 0,24 | 1,82 | 0,39 | 0,82 |
| 2 | *App*$^{NL-F}$ | | | | -1,16 | | 0,34 | -0,80 | 3,77 | -0,52 | | -0,41 | 1,49 | -0,52 | 0,18 |
| 3 | *App*$^{NL-F}$ | | | | -1,09 | -0,54 | | | | | | -0,74 | | | -0,74 |
| 3 | *App*$^{NL-F}$ | | | | -0,84 | 0,23 | | | | | | -0,03 | | | -0,03 |
| 3 | *App*$^{NL-F}$ | | | | -0,61 | -0,15 | | | | | | -0,38 | | | -0,38 |
| 3 | *App*$^{NL-F}$ | | | | -1,67 | -0,15 | | | | | | -0,40 | | | -0,40 |

***Table S1 : Calculation of the emotionality z-score*** *(related to figure 5)*

*For each test, a z-score ($z = (X - \mu)/\sigma$) is calculated for each parameter of consideration (zSC_param). All the zSC-parameters are then averaged per test leading to a test z-score (zSc_test). The average of each score gives the final z-score per animal.*

*X: individual values for the considered parameter; μ: mean of the control group; σ: standard deviation of the control group. In this figure, C57Bl6 mice have been used as the control group.*

*Table S2*

| | | zSc_param | | | | | | | | | | | | zSc_test | | | | |
|---|---|---|---|---|---|---|---|---|---|---|---|---|---|---|---|---|---|---|
| **Cohort** | **Treatment** | **zSC_OF_Distance** | **zSC_OF_Center entries** | **zSC_OF_Center time** | **zSC_OF_Center/Total distance ratio** | **zSC_EPM_Open arms entries** | **zSC_EPM_Open arms time** | **zSC_EPM_Open/Total arms entries** | **zSC_ST__Grooming latency** | **zSC_ST_Grooming frequency** | **zSC_ST_Grooming duration** | **zSC_TST_Immobility latency** | **zSC_TST_Immobility duration** | **zSC_OF** | **zSC_EPM** | **zSC_ST** | **zSC_TST** | **Emotionality z-score** |
| *Formula* | | *(X-μ)/σ* | | | | | | | | | | | | *Average parameter (per test)* | | | | *Average z-score* |
| 1 | $App^{NL-F}$ + NaCl | 0.61 | 0.45 | 0.33 | 0.21 | | | | | | | 1.59 | -0.22 | 0.40 | | | 0.68 | 0.54 |
| 1 | $App^{NL-F}$ + NaCl | 1.37 | 0.02 | -0.52 | -1.35 | | | | | | | 0.42 | 0.92 | -0.11 | | | 0.67 | 0.27 |
| 1 | $App^{NL-F}$ + NaCl | 0.29 | 0.56 | 0.25 | 0.30 | | | | | | | -1.06 | 0.77 | 0.35 | | | -0.14 | 0.10 |
| 1 | $App^{NL-F}$ + NaCl | 1.27 | 1.32 | 1.61 | 1.29 | | | | | | | 0.95 | -1.63 | 1.37 | | | -0.33 | 0.51 |
| 1 | $App^{NL-F}$ + NaCl | -0.23 | 0.23 | -1.87 | -1.29 | | | | | | | -0.42 | -0.48 | -0.79 | | | -0.45 | -0.62 |
| 1 | $App^{NL-F}$ + NaCl | -0.26 | 0.45 | -0.27 | 0.11 | | | | | | | 0.42 | -0.87 | 0.01 | | | -0.22 | -0.10 |
| 1 | $App^{NL-F}$ + NaCl | 1.48 | 0.88 | 1.15 | 0.63 | | | | | | | -1.06 | 0.18 | 1.04 | | | -0.43 | 0.30 |
| 1 | $App^{NL-F}$ + NaCl | 0.41 | -0.19 | -1.21 | -1.20 | | | | | | | -0.85 | 1.33 | -0.54 | | | 0.24 | -0.15 |
| 2 | $App^{NL-F}$ + NaCl | -0.50 | 0.23 | 1.23 | 1.48 | -0.99 | -1.78 | -1.52 | 0.03 | -1.32 | 1.32 | | | 0.61 | -1.43 | 0.01 | | -0.27 |
| 2 | $App^{NL-F}$ + NaCl | -0.12 | 0.67 | 0.88 | 1.06 | 0.77 | 0.44 | 0.79 | -0.94 | -1.32 | 1.39 | | | 0.62 | 0.67 | -0.28 | | 0.33 |
| 2 | $App^{NL-F}$ + NaCl | 1.60 | 1.21 | 1.58 | 1.11 | -0.11 | -2.28 | -0.25 | -0.26 | 1.32 | -0.06 | | | 1.38 | -0.88 | 0.33 | | 0.27 |
| 2 | $App^{NL-F}$ + NaCl | 0.66 | -0.19 | -0.97 | -1.17 | 0.77 | 0.44 | 0.79 | 1.09 | 0 | -0.32 | | | -0.42 | 0.67 | 0.25 | | 0.16 |
| 2 | $App^{NL-F}$ + NaCl | -0.60 | -1.06 | -0.26 | 0.27 | -0.11 | -1.93 | 0.04 | 0.55 | 0 | -0.95 | | | -0.41 | -0.66 | -0.13 | | -0.40 |
| 2 | $App^{NL-F}$ + NaCl | -1.13 | -2.47 | -1.52 | -0.44 | 0.77 | 0.44 | 0.79 | -1.47 | 0 | -1.47 | | | -1.39 | 0.67 | -0.98 | | -0.56 |
| 2 | $App^{NL-F}$ + NaCl | -0.36 | -0.84 | -1.92 | -1.24 | 0.77 | 0.44 | 0.79 | 1.46 | 1.32 | 0.24 | | | -1.09 | 0.67 | 1.01 | | 0.19 |
| 2 | $App^{NL-F}$ + NaCl | -1.19 | -1.28 | -0.70 | 0.18 | -1.88 | -1.70 | -1.44 | -0.46 | 0 | -0.14 | | | -0.74 | -1.67 | -0.20 | | -0.87 |
| 1 | $App^{NL-F}$ + ketamine | 0.41 | -0.41 | -4.72 | -4.66 | | | | | | | 1.27 | -0.45 | -2.34 | | | 0.40 | -0.96 |
| 1 | $App^{NL-F}$ + ketamine | -1.85 | -2.14 | -1.65 | -0.20 | | | | | | | -2.66 | 2.57 | -1.46 | | | -0.04 | -0.75 |
| 1 | $App^{NL-F}$ + ketamine | -0.08 | 0.56 | 0.35 | 0.58 | | | | | | | 0.85 | 1.01 | 0.35 | | | 0.93 | 0.64 |
| 1 | $App^{NL-F}$ + ketamine | -1.26 | -1.71 | -1.44 | -0.32 | | | | | | | 1.38 | 0.86 | -1.18 | | | 1.12 | -0.03 |
| 1 | $App^{NL-F}$ + ketamine | -0.94 | -1.06 | -1.16 | -0.27 | | | | | | | -3.29 | 1.21 | -0.86 | | | -1.04 | -0.95 |
| 1 | $App^{NL-F}$ + ketamine | 0.33 | -0.63 | -2.18 | -2.08 | | | | | | | -2.76 | 1.39 | -1.13 | | | -0.68 | -0.91 |
| 1 | $App^{NL-F}$ + ketamine | 0.60 | 0.02 | -1.06 | -1.20 | | | | | | | -2.34 | 1.42 | -0.41 | | | -0.45 | -0.43 |
| 1 | $App^{NL-F}$ + ketamine | -0.29 | -0.63 | -0.35 | 0.05 | | | | | | | -0.74 | 1.98 | -0.30 | | | 0.61 | 0.15 |
| 2 | $App^{NL-F}$ + ketamine | 3.42 | 1.75 | -0.76 | -6.51 | -0.11 | 0.16 | -0.94 | -2.35 | 2.64 | 5.55 | | | -0.52 | -0.29 | 1.94 | | 0.37 |
| 2 | $App^{NL-F}$ + ketamine | 1.43 | 0.67 | 0.83 | 0.26 | -0.99 | -4.40 | -1.52 | -1.17 | 2.64 | 4.68 | | | 0.79 | -2.30 | 2.05 | | 0.18 |
| 2 | $App^{NL-F}$ + ketamine | 0.64 | -0.30 | 0.63 | 0.50 | -0.11 | 0.27 | -0.69 | -1.77 | 2.64 | 2.96 | | | 0.36 | -0.17 | 1.28 | | 0.49 |
| 2 | $App^{NL-F}$ + ketamine | -0.13 | -0.08 | 0.50 | 0.74 | -0.99 | -1.56 | -1.52 | -1.82 | -1.32 | -0.70 | | | 0.25 | -1.36 | -1.28 | | -0.79 |
| 2 | $App^{NL-F}$ + ketamine | 1.75 | 1.21 | 2.04 | 1.67 | -0.99 | -3.68 | -1.29 | -2.06 | 5.29 | 3.55 | | | 1.67 | -1.99 | 2.26 | | 0.64 |
| 2 | $App^{NL-F}$ + ketamine | 0.46 | -0.19 | -0.37 | -0.40 | 0.77 | 0.44 | 0.79 | -0.42 | 0 | -0.57 | | | -0.12 | 0.67 | -0.33 | | 0.07 |
| 2 | $App^{NL-F}$ + ketamine | 1.09 | 1.10 | 0.75 | 0.39 | -0.11 | 0.02 | -0.07 | -0.73 | 3.96 | 2.04 | | | 0.83 | -0.05 | 1.75 | | 0.84 |
| 2 | $App^{NL-F}$ + ketamine | 0.69 | 0.45 | -2.91 | -3.21 | -0.11 | -0.26 | -0.36 | -2.02 | 6.61 | 2.70 | | | -1.24 | -0.24 | 2.43 | | 0.31 |

**Table S2 :** ***Calculation of the emotionality z-Score*** *(related to figures 6)*

*For each paradigm, a z-score ($z = (X - \mu)/\sigma$) is calculated for each parameter of consideration (zSC_param). All the zSC-parameters are then averaged per test leading to a test z-score (zSc_test). The average of each score gives the final z-score per animal.*

*X: individual values for the considered parameter; μ: mean of the control group; σ: standard deviation of the control group. In this figure, $App^{NL\text{-}F}$ + NaCl mice have been used as the control group.*

*Table S3*

| **Parameter** | **Statistical test** | **Statistical value** | **p-value** | **Significance** |
|---|---|---|---|---|
| *ELISA* | | | | |
| Membrane fraction | Two-way ANOVA | Structure: 12.335<br>Genotype: 23.529<br>Interact.: 8.039 | Structure: <0.001<br>Genotype: <0.001<br>Interact.: 0.002 | Structure: ***<br>Genotype: ***<br>Interact.: *** |
| Membrane fraction | Bonferroni multiple comparison test | | Cortex: 0.600<br>Hippocampus: <0.001<br>Cerebellum: 0.170 | Cortex: ns<br>Hippocampus: ***<br>Cerebellum: ns |
| Soluble fraction | Two-way ANOVA | Structure: 0.613<br>Genotype: 19.126<br>Interact.: 0.165 | Structure: 0.440<br>Genotype: <0.001<br>Interact.: 0.849 | Structure: ns<br>Genotype: ***<br>Interact.: ns |
| Soluble fraction | Bonferroni multiple comparison test | | Cortex: 0.584<br>Hippocampus: 0.938<br>Cerebellum: 0.513 | Cortex: ns<br>Hippocampus: ns<br>Cerebellum: ns |
| Aggregates | Two-way ANOVA | Structure: 17.935<br>Genotype: 24.665<br>Interact.: 0.887 | Structure: <0.001<br>Genotype: <0.001<br>Interact.: 0.423 | Structure: ***<br>Genotype: ***<br>Interact.: ns |
| Aggregates | Bonferroni multiple comparison test | | Cortex: 0.008<br>Hippocampus: 0.004<br>Cerebellum: 0.101 | Cortex: **<br>Hippocampus: **<br>Cerebellum: ns |
| *ASTROCYTE MORPHOLOGY* | | | | |
| Sholl analysis | Two-way ANOVA | Distance to soma: 98.167<br>Genotype: 26.845<br>Interact.: 19.727 | Structure: <0.001<br>Genotype: <0.001<br>Interact.: <0.001 | Structure: ***<br>Genotype: ***<br>Interact.: *** |
| Sholl analysis | Mann-Whitney multiple comparison test | | 5 µm: 0.049<br>10µm: <0.001<br>15µm: <0.001<br>20µm: <0.001<br>25µm: <0.001<br>30µm: 0.003<br>35µm: 0.214<br>40µm: 0.121<br>45µm: 0.172 | 5 µm: *<br>10µm: ***<br>15µm: ***<br>20µm: ***<br>25µm: ***<br>30µm: **<br>35µm: ns<br>40µm: ns<br>45µm: ns |
| Number of branches | Mann-Whitney U test | 411 | <0.001 | *** |
| Average cellular volume | Mann-Whitney U test | 74.5 | 0.001 | ** |
| *ELECTROPHYSIOLOGY* | | | | |
| Tonic GABA | Mann-Whitney U test | 41 | 0.038 | * |
| Patch_RMP | Student t-test | -2.326 | 0.032 | * |
| Patch_Frequency sEPSC | Student t-test | 1.177 | 0.255 | ns |
| Patch_Amplitude sEPSC | Student t-test | 0.302 | 0.766 | ns |
| Patch_Rise sEPSC | Mann-Whitney U test | 46.500 | 0.896 | ns |
| Patch_Decay sEPSC | Student t-test | 0.144 | 0.887 | ns |
| Patch_Frequency mEPSC | Student t-test | 3.087 | 0.006 | ** |
| Patch_Amplitude mEPSC | Student t-test | -1.001 | 0.330 | ns |
| Patch_Rise mEPSC | Mann-Whitney U test | 40 | 0.481 | ns |
| Patch_Decay mEPSC | Student t-test | 0.412 | 0.685 | ns |
| fEPSP vs. 1 at t0 | Student t-test | C57Bl6 controls: 6.805<br>AppNL-F: 6.868 | C57Bl6 controls: <0.001<br>AppNL-F: <0.001 | C57Bl6 controls: ***<br>AppNL-F: *** |

| | | | | |
|---|---|---|---|---|
| fEPSP_t1-5min | Student t-test | 2.629 | 0.01 | * |
| fEPSP vs. 1 at t55 | Student t-test | C57Bl6 controls: 3.521<br>AppNL-F: 7.205 | C57Bl6 controls: 0.001<br>AppNL-F: <0.001 | C57Bl6 controls: **<br>AppNL-F: *** |
| fEPSP_t55-60min | Student t-test | 15.926 | <0.001 | *** |
| *ELECTROPHYSIOLOGY ± DEPRENYL* | | | | |
| Patch_RMP | Student t-test | 2.238 | 0.042 | * |
| Patch_Frequency sEPSC | Mann-Whitney U test | 48 | 0.105 | Ns |
| Patch_Amplitude sEPSC | Student t-test | 1.883 | 0.012 | ** |
| Patch_Rise sEPSC | Mann-Whitney U test | 25 | 0.505 | ns |
| Patch_Decay sEPSC | Student t-test | -0.387 | 0.705 | ns |
| Patch_Frequency mEPSC | Mann-Whitney U test | 95 | 0.003 | ** |
| Patch_Amplitude mEPSC | Student t-test | 2.838 | 0.013 | * |
| Patch_Rise mEPSC | Student t-test | 0.317 | 0.756 | ns |
| Patch_Decay mEPSC | Mann-Whitney U test | 22 | 0.328 | ns |
| fEPSP vs. 1 at t0 | Student t-test | $App^{NL-F}$ + aCSF: 204<br>$App^{NL-F}$ + deprenyl: 465 | $App^{NL-F}$ + aCSF: 0.004<br>$App^{NL-F}$ + deprenyl: <0.001 | $App^{NL-F}$ + aCSF: **<br>$App^{NL-F}$ + deprenyl: *** |
| fEPSP_t1-5min | Mann-Whitney test | 204 | 0.004 | ** |
| fEPSP vs. 1 at t55 | Student t-test | $App^{NL-F}$ + aCSF: 139<br>$App^{NL-F}$ + deprenyl: 435 | $App^{NL-F}$ + aCSF: <0.001<br>$App^{NL-F}$ + deprenyl: <0.001 | $App^{NL-F}$ + aCSF: ***<br>$App^{NL-F}$ + deprenyl: *** |
| fEPSP_t55-60min | Mann-Whitney U test | *325* | *<0.001* | *** |
| *BEHAVIOR* | | | | |
| OF_distance | Mann-Whitney U test | 4.702 | <0.001 | *** |
| OF_Center entries | Mann-Whitney U test | 3.397 | 0.008 | ** |
| OF_Center time | Mann-Whitney U test | -0.062 | 0.952 | ns |
| EPM_Open arm entries | Student t-test | 2.602 | 0.021 | * |
| EPM_Open arms time | Student t-test | 1.451 | 0.169 | ns |
| FST_Immobility latency | Mann-Whitney U test | 83.5 | 0.036 | * |
| TST_Immobility duration | Student t-test | -3.994 | <0.001 | *** |
| SP_Grooming latency | Student t-test | 1.440 | 0.162 | ns |
| SP_Grooming frequency | Student t-test | -0.163 | 0.874 | ns |
| SP_Grooming duration | Mann-Whitney U test | 112 | 0.325 | ns |
| SPT_Sucrose consumption | Student t-test | -0.112 | 0.912 | ns |
| Emotionality z-score | Student t-test | -2.770 | 0.009 | ** |
| 5-trial social memory test_Latency | One-way ANOVA | Tiral: 6.057<br>Genotype: 5.463<br>Interact.: 10.548 | Trial: 0.027<br>Genotye: 0.035<br>Interact.: 0.006 | Trial: *<br>Genotype: *<br>Interact.: ** |
| 5-trial social memory test_Latency | Bonferroni multiple comparison test | | Tiral 1 vs. Trial 5<br>C57Bl6 controls: >0.999<br>AppNL-F: 0.0037<br><br>Genotype<br>Trial 1: 0.872<br>Tiral 5: 0.001 | Tiral 1 vs. Trial 5<br>C57Bl6 controls: ns<br>AppNL-F: **<br><br>Genotype<br>Trial 1: ns<br>Tiral 5: ** |
| 5-trial social memory test_Interactions | One-way ANOVA | Tiral: 25.915<br>Genotype: 0.014<br>Interact.: 5.367 | Trial: <0.001<br>Genotye: 0.908<br>Interact.: 0.007 | Trial: ***<br>Genotype: ns<br>Interact.: *** |
| 5-trial social memory test_Interactions | Bonferroni multiple comparison test | | C57Bl6 controls<br>1 vs. 2: 0.005<br>1 vs. 3: 0.003<br>1 vs. 4: 0.047<br>1 vs. 5: >0.999<br>4 vs. 5: 0.039 | C57Bl6 controls<br>1 vs. 2: **<br>1 vs. 3: **<br>1 vs. 4: *<br>1 vs. 5: ns<br>4 vs. 5: * |

| | | | | |
|---|---|---|---|---|
| | | | AppNL-F<br>1 vs. 2: 0.002<br>1 vs. 3: <0.001<br>1 vs. 4: 0.004<br>1 vs. 5: 0.119<br>4 vs. 5: >0.999 | AppNL-F<br>1 vs. 2: **<br>1 vs. 3: ***<br>1 vs. 4: **<br>1 vs. 5: ns<br>4 vs. 5: ns |
| Olfactory habituation/deshavituation | One-way ANOVA | Odor: 6.114<br>Genotype: 0.018<br>Interact.: 0.029 | Odor: 0.009<br>Genotype: 0.893<br>Interact.: 0.953 | Odor: **<br>Genotype: ns<br>Interact.: ns |
| Olfactory habituation/deshavituation | Bonferroni multiple comparison test | | Melon vs. Banana: >0.999<br>Melon vs. Famil.: >0.999<br>Melon vs. Non famil.: 0.125<br>Banana vs. Famil.: 0.763<br>Banana vs. Non famil.: 0.035<br>Famil. Vs. Non famil.: 0.163 | Melon vs. Banana: ns<br>Melon vs. Famil.: ns<br>Melon vs. Non famil.: ns<br>Banana vs. Famil.: ns<br>Banana vs. Non famil.: *<br>Famil. Vs. Non famil.: ns |
| *BEHAVIOR ± KETAMINE* | | | | |
| OF_distance | Student t-test | -0.416 | 0.681 | ns |
| OF_Center entries | Student t-test | -0.728 | 0.473 | ns |
| OF_Center time | Student t-test | -1.461 | 0.156 | ns |
| EPM_Open arm entries | Mann-Whitney U test | 41.5 | 0.328 | ns |
| EPM_Open arms time | Mann-Whitney U test | 38 | 0.574 | ns |
| TST_Immobility latency | Mann-Whitney U test | -1.326 | 0.107 | ns |
| TST_Immobility duration | Student t-test | 2.651 | 0.019 | * |
| SP_Grooming latency | Student t-test | 3.603 | 0.003 | ** |
| SP_Grooming frequency | Student t-test | -2.862 | 0.013 | * |
| SP_Grooming duration | Student t-test | -2.905 | 0.012 | * |
| Emotionality z-score | Mann-Whitney U test | 125 | 0.926 | ns |
| 5-trial social memory test_Latency | Wilcoxon test | 2.249 | 0.025 | * |
| 5-trial social memory test_Latency | Miultiple comparison | | *Trial 1 vs. Trial 5<br>AppNL-F + NaCl: 0.111<br>AppNL-F + Ket.: 0.106* | *Trial 1 vs. Trial 5<br>AppNL-F + NaCl: ns<br>AppNL-F + Ket.: ns* |
| 5-trial social memory test_Interactions | Friedman ANOVA | 43.116 | <0.001 | *** |
| 5-trial social memory test_Interactions | Bonferroni multiple comparison test | | *AppNL-F + NaCl*<br>1 vs. 2: 0.219<br>1 vs. 3: <0.001<br>1 vs. 4: <0.001<br>1 vs. 5: <0.001<br>4 vs. 5: 0.442<br><br>AppNL-F + Ketamine<br>1 vs. 2: 0.710<br>1 vs. 3: 0.495<br>1 vs. 4: 0.022<br>1 vs. 5: 0.049<br>4 vs. 5: 0.710 | AppNL-F + NaCl<br>1 vs. 2: ns<br>1 vs. 3: ***<br>1 vs. 4: ***<br>1 vs. 5: ***<br>4 vs. 5: ns<br><br>AppNL-F + Ketamine<br>1 vs. 2: ns<br>1 vs. 3: ns<br>1 vs. 4: *<br>1 vs. 5: *<br>4 vs. 5: ns |
| ELECTROPHYSIOLOGY ± KETAMINE | | | | |
| Patch_RMP | Student t-test | -0.307 | 0.765 | ns |
| Patch_Frequency sEPSC | Mann-Whitney U test | 22 | >0.999 | ns |
| Patch_Amplitude sEPSC | Mann-Whitney U test | -1.960 | 0.083 | ns |
| Patch_Rise sEPSC | Mann-Whitney U test | -1.101 | 0.312 | ns |
| Patch_Decay sEPSC | Student t-test | 22 | >0.999 | ns |
| Patch_Frequency mEPSC | Student t-test | -4.655 | 0.008 | ** |

| | | | | |
|---|---|---|---|---|
| Patch_Amplitude mEPSC | Student t-test | -1.237 | 0.242 | ns |
| Patch_Rise mEPSC | Mann-Whitney U test | 21 | >0.999 | ns |
| Patch_Decay mEPSC | Student t-test | -1.328 | 0.235 | ns |
| fEPSP vs. 1 at t0 | Student t-test | AppNL-F + NaCl: 12.93<br>AppNL-F + Ket.: 2.835 | AppNL-F + NaCl: <0.001<br>AppNL-F + Ket.: 0.009 | AppNL-F + NaCl: ***<br>AppNL-F + Ket.: ** |
| fEPSP_t1-5min | Mann-Whitney U test | 189 | 0.168 | ns |
| fEPSP vs. 1 at t55 | Student t-test | AppNL-F + NaCl: 10.49<br>AppNL-F + Ket.: 3.626 | AppNL-F + NaCl: <0.001<br>AppNL-F + Ket.: 0.001 | AppNL-F + NaCl: ***<br>AppNL-F + Ket.: ** |
| fEPSP_t55-60min | Mann-Whitney U test | 208 | <0.001 | *** |
| Tonic GABA | Mann-Whitney U test | 45 | <0.001 | *** |

**Table 3:** ***Detailed statistics***

*A normality test and a Levene’s equality of variance test were performed to determine the subsequent analysis. 0.05 was set as the significance threshold.*
